# Cell-ID: gene signature extraction and cell identity recognition at individual cell level

**DOI:** 10.1101/2020.07.23.215525

**Authors:** Cortal Akira, Martignetti Loredana, Six Emmanuelle, Rausell Antonio

**Affiliations:** Paris University, Imagine Institute, 75015 Paris, France, EU; Clinical Bioinformatics Laboratory, INSERM UMR1163, Necker Hospital for Sick Children, 75015 Paris, France, EU; Laboratory of Human Lymphohematopoiesis, INSERM UMR 1163, Paris, France

## Abstract

The exhaustive exploration of human cell heterogeneity requires the unbiased identification of molecular signatures that can serve as unique cell identity cards for every cell in the body. However, the stochasticity associated with high-throughput single-cell sequencing has made it necessary to use clustering-based computational approaches in which the characterization of cell-type heterogeneity is performed at cell-subpopulation level rather than at full single-cell resolution. We present here Cell-ID, a clustering-free multivariate statistical method for the robust extraction of per-cell gene signatures from single-cell sequencing data. Cell-ID signatures allow unbiased cell identity recognition across different donors, tissues-of-origin, model organisms and single-cell omics technologies. Cell-ID is distributed as an open-source R software package: https://github.com/RausellLab/CelliD.

---

High-throughput single-cell technologies, such as single-cell RNA-seq, are currently being used for the complete cellular characterization of human tissues and cell types. The Human Cell Atlas project^1^, the NIH Human BioMolecular Atlas Program (HuBMAP)^2^ and the LifeTime^3^ initiative are remarkable examples of current scientific ambitions in this direction. One of the major goals of these studies is the identification of previously unknown cell types or cell states with potential physiological roles in health and disease. However, the computational characterization of cell heterogeneity is rendered more complex by the high dimensionality and high levels of technical and biological noise associated with single-cell measurements^4^. One common strategy for enhancing the signal-to-noise ratio involves the use of a low-dimensional representation of cells from which the most salient relative differences may emerge^5^. The techniques most widely used for this purpose include principal component analysis (PCA), independent component analysis (ICA), t-stochastic nuclear embedding (t-SNE) and uniform manifold approximation and projection (UMAP)^6^. Molecular characterization of the observed cell heterogeneity is then typically performed by analysing differential gene expression between the groups of cells resulting from clustering in low-dimensional space. However, this reliance on cell clusters results in gene signatures being assigned at cell subpopulation level rather than at full single-cell resolution. One of the major limitations of cluster-based approaches is that the gene signature analysis is bound to a certain level of resolution, at which the cell heterogeneity is partitioned into non-overlapping classes^7^. An exhaustive exploration of transcriptional heterogeneity requires a statistically robust per-cell gene signature assessment for each cell in a data set. No such approach has been described to date.

We present here Cell-ID, a multivariate approach to extracting a gene signature for each individual cell in a study (**Fig. 1** and **Online Methods**). Cell-ID is based on multiple correspondence analysis (MCA), a statistical technique based on singular-value decomposition (SVD) that can provide simultaneous projections of individuals (e.g. cells) and variables (e.g. genes) in the same low-dimensional space^8–10^. This represents a major advantage over alternative low-dimensional transformations providing only cell projections. Originally MCA applies to binary and fuzzy-coded data, and this has been so far an obstacle for its use on *omics* data. We describe here a linear scaling of gene expression values unlocking the use of this technique for quantitative single-cell data analysis, while keeping the MCA mathematical properties. By representing cells and genes on the same principal axes, analytical distances can be calculated not only between cells and between genes, but also between each cell and each gene (**Fig. 1A**). In such a space, the closer a gene *g* is to a cell *c*, the more specific to such a cell it can be considered. Gene-to-cell distances can then be ranked for each individual cell, and the top-ranked genes may be regarded as a unique gene signature representing the identity card of the cell (**Fig. 1A**). We show here that the unbiased per-cell gene signatures extracted by Cell-ID reproduce the gene signatures previously established for well-known cell types. Cell-ID signatures are, in turn, reproducible across independent datasets, despite strong batch effects, and can be used for cell identity tracing across different donors, model organisms, tissues-of-origin and single-cell omics protocols.

**Figure 1.**
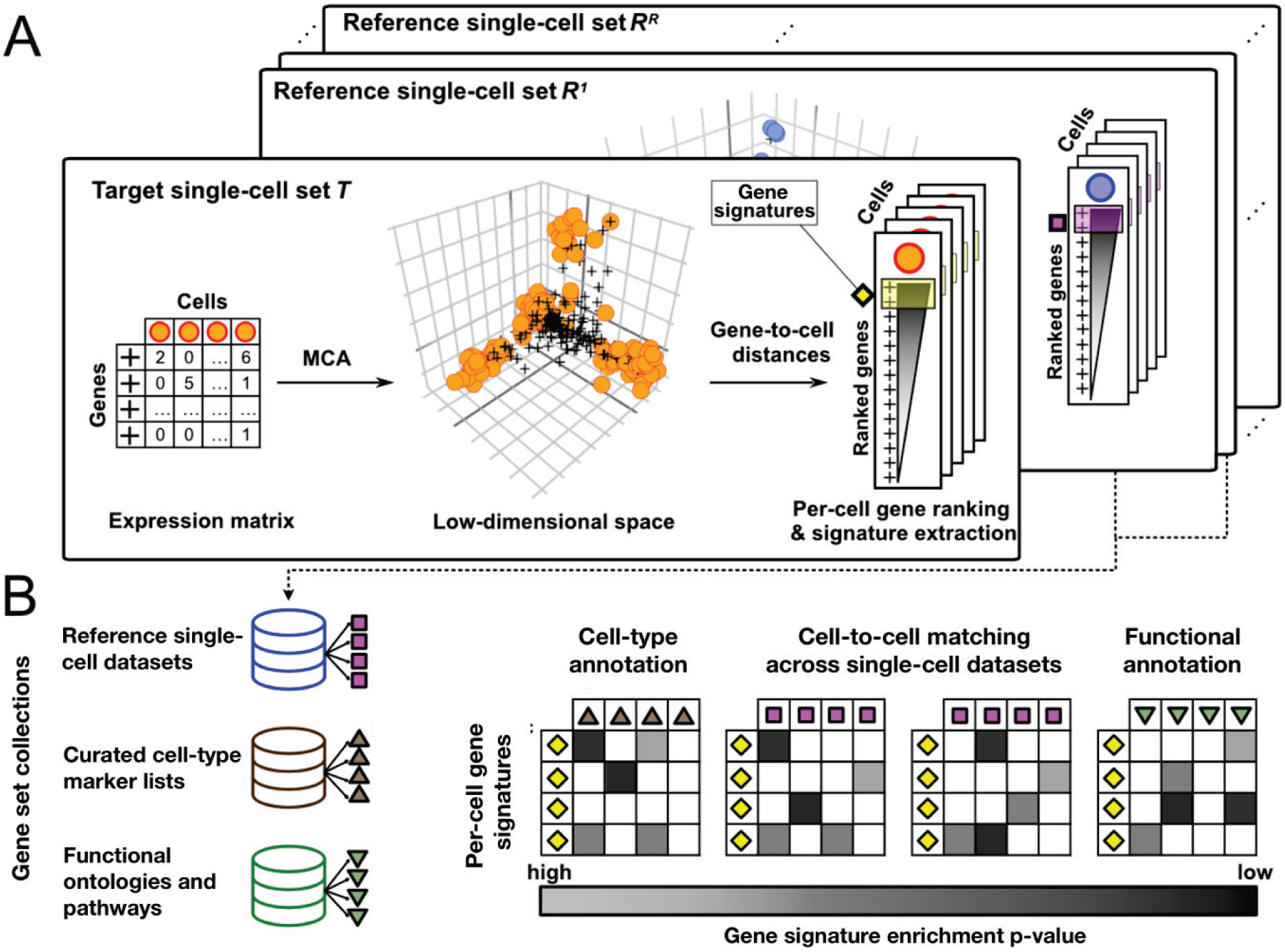
Overview of the Cell-ID approach. **(A)** Cell-ID performs a dimensionality reduction of the gene expression matrix through multiple correspondence analysis (MCA), where both cells and genes are projected in a common orthogonal space. In such space, the closer a gene is to a cell the more specific it is to it. Thus, a gene ranking is obtained for each cell in a dataset based on their distance in the MCA space. The top-ranked genes for a given cell define its gene signature, which can be regarded as a unique cell identity card. Per-cell gene signatures can be independently extracted for a collection of single-cell datasets for downstream analyses. **(B)** Per-cell gene signatures from a dataset can be evaluated for their enrichment against (i) collections of pre-established cell type markers, in order to perform automatic cell type annotation, (ii) per-cell gene signatures from independent single-cell datasets, allowing cell matching and label transferring, and (iii) gene sets representing biological functions or molecular pathways, allowing functional annotation and interpretation of cell states.

We first evaluated the consistency of MCA-based low-dimensional representations of cells and genes on 100 simulated single-cell RNA-seq datasets (**Supplementary Note 1**). Consistency was demonstrated at three levels. The cell representation achieved by MCA dimensionality reduction was largely equivalent to that achieved by principal component analysis (PCA) on the same dataset: Spearman’s correlation coefficients, for the correlation between MCA and PCA coordinates, with median and standard deviation values across data sets of 1.00±0.02, 1.00±0.02, 0.99±0.02, 0.99±0.02, and 0.99±0.02 for their first five principal axes, respectively (**Supplementary Fig.1** and **Supplementary Note 1**). Second, the per-cell gene rankings provided by MCA are consistent with the expression values for the 50 neighbouring cells in MCA space, as reflected by their log-fold change in expression relative to the other cells (**Supplementary Fig. 2A and 2B**). Third, MCA-based per-cell gene rankings are robust to high levels of dropout events. Thus, genes with no expression detected in a cell may nevertheless rank highly for the cell concerned if more strongly expressed in the surrounding cells than in more distant cells (**Supplementary Fig. 2A** and **2B**; **Supplementary Note 1)**. Our results highlight the advantages of a multivariate approach in which per-cell gene assessments are implicitly weighted by their cell neighbourhood in low-dimensional space. Such patterns would be missed if per-cell gene rankings were naively obtained either (i) from the log-fold changes in expression in the target cell relative to background cells (**Supplementary Fig. 2C-D)** or (ii) from highest-to-lowest expression values within a cell, with random ranks for ties, as in another published approach (AUCell^11^, SCINA^12^) (**Supplementary Fig. 2E-F**).

We also showed that Cell-ID could extract per-cell gene signatures, recovering characteristic marker lists associated with well-known cell types^13^ (**Supplementary Note 2**). To this end, we used two independent sets of human blood mononuclear cells for which individual cells were confidently annotated with an actual cell type through concomitant measurements of single-cell protein marker levels: (i) cord blood mononuclear cells (CBMCs) profiled with a CITE-seq protocol^14^; and (ii) peripheral blood mononuclear cells (PBMCs) profiled with a REAP-seq protocol^15^. Cell-ID per-cell gene signatures were significantly enriched in the lists of markers associated with the corresponding cell type (**Figure 2**). This enrichment made it possible to recognize cell types with high rates of precision (87% and 90%) and recall (84% and 73%) for both datasets (multinomial *p*-value < 10^−16^ for all figures), outperforming reference methods for cell-type classification on the basis of marker lists (AUCELL^11^ and SCINA^12^; **Fig. 2B, Supplementary Fig. 3-4**). In more challenging scenarios, Cell-ID was capable of non-disjoint multi-class cell-type assignments capturing smooth transitions between hematopoietic differentiation states from the most immature hematopoietic stem cell (HSC) to downstream myeloid (CMP/GMP) and erythroid progenitors (MEP) (**Fig. 2, C-D**). It was consistently able to identify singleton cells, i.e. rare cell types represented by only one cell within a dataset (**Supplementary Note 2)**. The capacity of Cell-ID to recover well-established cell types at single-cell resolution supports its use for automated cell-type annotation, even for extremely rare cells, without the need for clustering.

**Figure 2.**
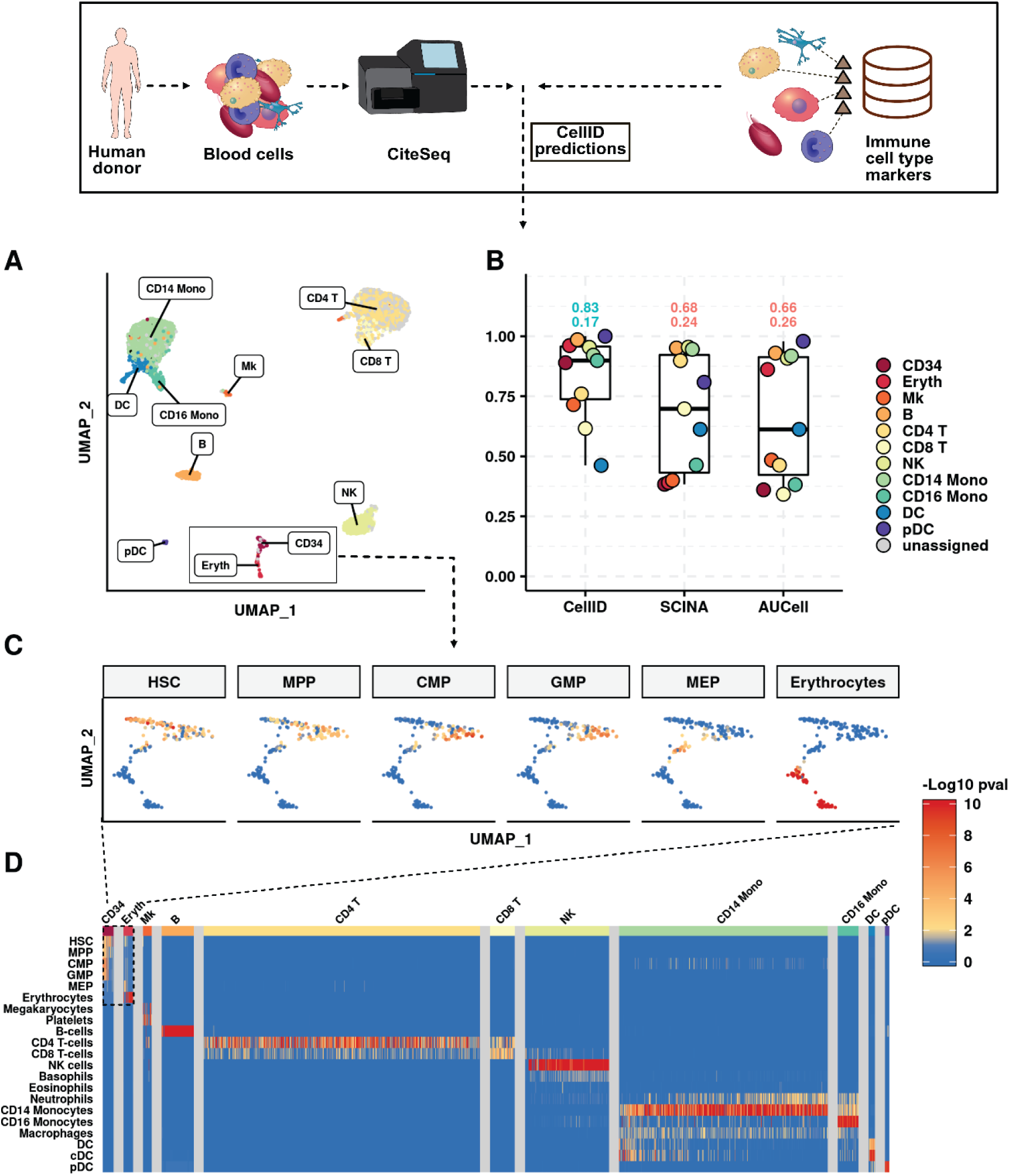
Cell-ID cell type prediction of human cord blood mononuclear cells using pre-established marker lists. (**A**) UMAP representation of 8005 cord blood mononuclear cells profiled by CITE-seq^3^. Dots representing cells are colored according to Cell-ID cell type predictions using pre-established immune cell signatures, as indicated by the labels in the figure. (**B**) Performance measured through the F1 score achieved by Cell-ID, AUCell and SCINA cell type predictions for each of the blood cell types reported in the original publication^3^. Boxplots summarize the F1 scores for each method. The numbers above boxplots denote the global performance (macro F1 score, upper digits) and its standard deviation (lower digits), where the maximum and minimum values across methods are colored in black and grey, respectively. (**C**) Zoomed UMAP representation on Erythrocytes and CD34+ cells showing that the Cell-ID multi-class cell assignments capture transient cell states consistent with the cell-type hierarchy associated with immature hematopoietic stem cell differentiation. Cells are color-coded according to the −log10 enrichment p-value obtained by Cell-ID in tests of the association of their gene signature with the cell-type signatures associated to their pre-cursor cell types: HSC, MPP, CMP, GMP, MEP and erythrocytes. The color scale for cells extends from white (*p* value=1) to dark red (*p* value = 1e-10), with p-values<1e-10 fixed at this value). **D)** Heatmap representing, for each individual cell (displayed in columns), the −log10 transformed *p*-value obtained by Cell-ID in tests of the association of the gene signature with each of the evaluated pre-established marker lists, representing a total of 21 blood cell types (displayed in rows). The color code of the heatmap extends from dark blue (*p* value=1) to yellow (*p* value = 10^−2^) to dark red (*p* value = 10^−10^), with *p* values<10^−10^ fixed at this value). Non-significant associations (*p* value>10^−2^ after Benjamini Hochberg correction for the number of gene signatures tested) are shown in blue. The columns in the heatmaps were grouped by the reference cell-type label, as indicated by the colored bands at the top and in the associated legend. CD34: CD34+ hematopoietic stem cells; Eryth: Erythrocytes; Mk: Megakaryocytes, B: B cells, CD4 T: CD4+ T cells, CD8: CD8+ T cells, CD14: CD14+ monocytes, CD16 Mono: CD16+ monocytes, NK: natural killer cells, DC: denderitic cells, pDC: plasmacytoid dendritic cell, HSC: hematopoietic stem cells, MPP: muti-potent-progenitor, CMP: common-myeloid-progenitors, GMP: granulocyte-monocyte-progenitor, MEP: megakryocyte-erythrocyte-progenitor.

We benchmarked the capacity of Cell-ID to match cells of analogous cell types across independent single-cell RNA-seq datasets from the same tissue-of-origin, within and across species (**Supplementary Note 3**). Cell-ID matching across datasets is performed by a per-cell assessment in the *query* dataset evaluating the replication of gene signatures extracted from the *reference* dataset. Gene signatures from the *reference* dataset can be automatically derived either from individual cells (Cell-ID(c)), or from previously-established groups of cells (Cell-ID(g); **Methods**). We thus analysed independent human pancreatic islets^16–18^ and human and mouse airway epithelium datasets^19,20^ corresponding to multiple donors and diverse sequencing technologies (**Fig. 3** and **Supplementary Note 3**). Cell-ID(c) and Cell-ID(g) consistently yielded high precision (>94% and >92%), recall (>77% and >75%) and F1 values (>88% and >87%, respectively) across all evaluated *reference*-to-*query* cell-type assignments (multinomial *p*-value < 2.2e-16; **Fig. 3A** and **Supplementary Fig. 5**). The overall performance of Cell-ID was at least as good as that of reference methods for cell matching and label transfer^21–29^ (**Fig. 3A, Supplementary Fig. 6, A-B**), and salient scores were obtained for cell types observed at low frequencies (<2%): epsilon cells, tissue-resident macrophages, mast cells and endothelial cells from pancreatic islet samples^17,18^, and PNEC, brush cells and ionocytes in mouse and human airway epithelium datasets^19,20^ (**Fig. 3B, Supplementary Fig. 6, C-D**). We further probed the capacity of this tool at individual cell resolution, by extracting Cell-ID signatures from 13 Schwann cells described in a dataset of 8,629 human pancreatic cells^16^. With these signatures, we were able to recognize four and two prototypic Schwann cells that had previously gone unnoticed within two independent sets of 2126 and 2261 human pancreatic cells, respectively, obtained from different donors and with different sequencing protocols^17,18^ (**Supplementary Figure 7**).

**Figure 3.**
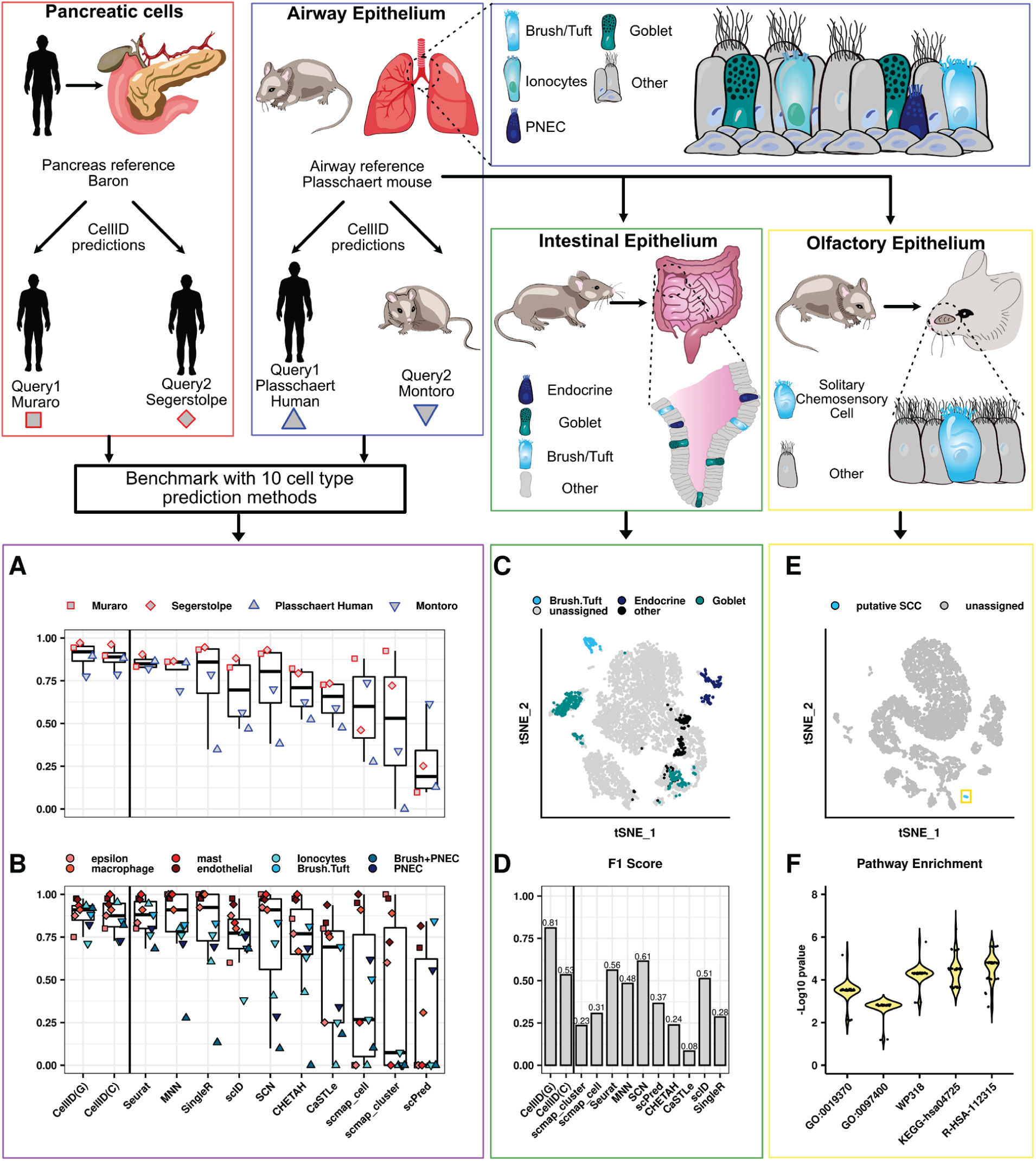
Performance of Cell-ID cell-to-cell matching across independent scRNA-seq datasets from the same or different tissue of origin, within and across species. **(A** and **B)** Performance measured through the F1 score (y-axis) achieved by Cell-ID(g), Cell-ID(c) and 10 alternative state-of-the-art methods (x-axis), covering the major approaches for cell-matching or label transfer across scRNA-seq datasets (**Supplementary Table 9**). **(A)** The performance for each method is represented for each of the label transferring evaluated (as schematically represented in the top left panels), corresponding to cell-to-cell matching across datasets from pancreatic islets cells (red squares and red diamonds), and across datasets from airway epithelium (blue triangles). Boxplots summarize the global performance (macro F1 scores) for each method. **(B)** The performance for each method is represented for each of the rare cell types reported in the original publications associated to the pancreatic islets cells (squares and diamonds) and airway epithelium datasets (blue triangles) evaluated in **(A)**. Rare cell types are represented following the color palette indicated in the legend and representing epsilon, macrophages, mast, and endothelial cells (pancreatic cells), and ionocytes, brush/tuft, and PNEC (airway epithelium. A cell type label gathering together brush and PNEC cells was used for consistency with the labelling provided in the human sample from Plasschaert et. al. (**C**) t-SNE representation of 7216 cells from mouse small intestinal epithelium ^29^. Dots representing cells are colored according to Cell-ID(g) cell type predictions, using as a reference the mouse airway epithelial gene signatures extracted from Plasschaert et al, as schematically represented in the panels above. Intestinal epithelium cell types with significant enrichment p-values are colored in green (goblet), brush/tuft (light blue) and dark blue (endocrine). Cells with significant enrichments in airway epithelial signatures without an analogous cell type in intestinal epithelium are represented in black. Intestinal epithelium cells with no significant enrichments were left unassigned and are displayed in grey. For comparison, the associated manual cell type annotations provided in ^29^ are represented in **Supplementary Figure 10 A**. (**D**) F1 score (y-axis) achieved by Cell-ID(g), Cell-ID(c) and 10 alternative state-of-the-art methods (x-axis), for the label transferring depicted in **h**. (**E**) UMAP representation of 9126 mouse olfactory epithelium cells from ^30^. Dots representing cells are colored according to Cell-ID(g) cell type predictions using as a reference the mouse brush/tuft gene signatures extracted from (i) mouse airway epithelium, and (ii) mouse small intestinal epithelium, as schematically represented in the panels above. The 37 cells significantly enriched with airway Brush/Tuft gene signatures are highlighted in blue and were interpreted as putative SCCs. Identical results were obtained when using intestinal Brush/Tuft gene signatures. (**F**) Violin plots corresponding to the distribution of −log10 enrichment p-values (y-axis) across the 37 cells identified in **(E)** (black dots) are represented for 5 significant functional terms (x-axis). GO-0019370: Gene Ontology term “leukotriene biosynthetic process”; GO-0097400: Gene Ontology term “interleukin-17-mediated signalling pathway”; WP318: WikiPathways term “Eicosanoid Synthesis”; KEGG-hsa04725: KEGG term “Cholinergic synapse”; R-HSA-112315: Reactome term “Transmission across Chemical Synapses”.

We then assessed the ability of Cell-ID to recognize rare cells from the same cell type across different tissues, and, thus, within diverse cell composition contexts (**Supplementary Note 4**). Thus, based on the unbiased gene signatures obtained from airway epithelium cells^19^, Cell-ID was able to identify brush/tuft cells, endocrine cells and goblet cells in the intestinal epithelium^30^ with high precision (92%), recall (73%) and F1 scores (81%), outperforming reference methods for cell matching (multinomial *p*-value < 2.2e-16 for all figures; **Fig. 3 C-D, Supplementary Fig. 8)**. From a discovery perspective, we used Cell-ID to perform cell-type scanning of two independent olfactory epithelium datasets ^31 32^ against brush/tuft signatures from the airway and the intestinal epithelium, which enabled us to identify putative rare solitary chemosensory cells (SCCs), a type of chemosensory cells closely related to brush/tuft cells, that had remained unclassified in the original publications. (**Fig. 3 E-F, Supplementary Fig. 9**). Thus, a total of 37 (<0.5%) and 5 (<0.6%) olfactory epithelium cells were found to display significant enrichment in airway and intestinal brush/tuft signatures (BH corrected *p* value< 10e-20), with a median of 29%±0.5 and 23%±0.4 of genes, respectively, in common. These cells displayed high levels of expression for the characteristic SCC marker genes *Il25* and *Gnat3*, and their Cell-ID gene signatures were significantly enriched in cysteinyl leukotriene biosynthesis genes, as reported by Ualiyeva et al^33^ for SCCs (**Supplementary Table 1**). Our findings confirm, at single-cell resolution, the recently reported transcriptional and functional similarity between rare olfactory SCCs and brush/tuft cells from the airways and intestinal epithelia^33,34^ (**Supplementary Note 4**).

Finally, we assessed the reproducibility of Cell-ID gene signatures across datasets profiled with different single-cell omics technologies: single-cell RNA-seq from the Tabula Muris mouse cell atlas^35^ and single-cell ATAC-seq from the Mouse ATAC Atlas^36^ (**Supplementary Note 5**). We benchmarked large-scale cell-type label transfer between the two expert-annotated atlases, collectively including 50 cell types from the eight tissues common to both: heart, kidney, liver, lung, bone marrow, spleen, thymus and large intestine (**Fig4. A-B, Supplementary Fig. 10**). Both Cell-ID(c) and Cell-ID(g) matched cell types across scRNA-seq and sciATAC-seq datasets with high F1 scores and, together with SingleR^28^, outperformed all the other reference methods evaluated (**Fig4. C-D, Supplementary Fig 11, Supplementary Figure 12)**. The capacity of Cell-ID to extract **gene signatures** that are robustly replicated across different single-cell *omics* technologies, in an automated manner, and its computational scalability (**Supplementary Note 6**) pave the way for the systematic multi-omics scanning of rare cell types across tissues and whole organisms.

**Figure 4.**
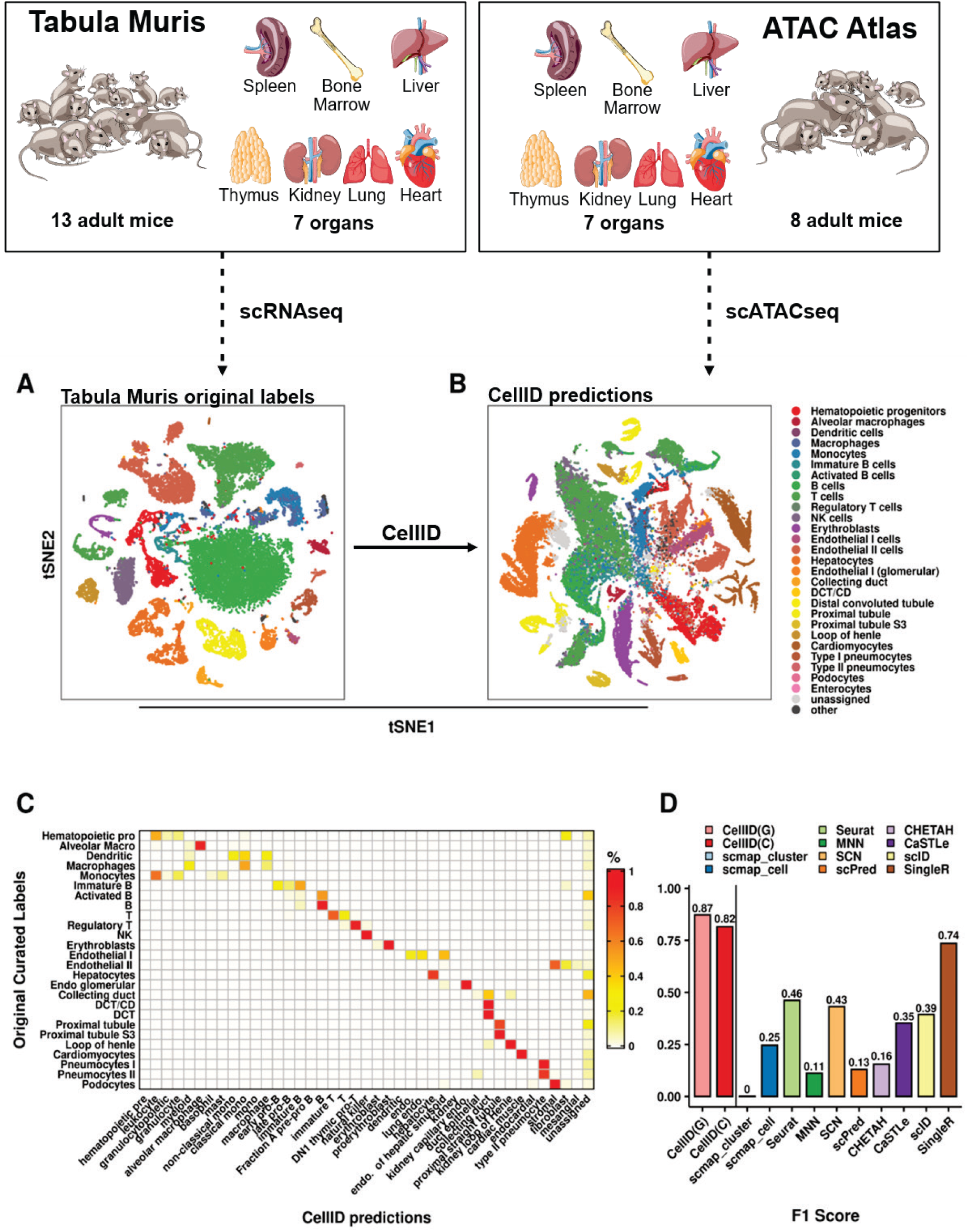
Performance of Cell-ID cell-to-cell matching across independent datasets from different single-cell *omics* technologies: scRNAseq and scATACseq. **(A)** t-SNE representation of 25332 cells from 7 organs profiled with scRNAseq 10X genomics from the Tabula Muris mouse cell atlas. Cells are colored according to the manually annotated cell types provided by the atlas, regrouped by the categories represented in the legend in **(B)** and described in **Supplementary Table 10. (B)** t-SNE representation of 50284 cells from 7 organs profiled with scATAC-seq from the Mouse ATAC atlas. Cells are colored according to the Cell-ID(g) predictions using the group gene signatures extracted from **(A)**, as represented in the legend. (**C**) Heatmap representing the confusion matrix between the manually curated cell type annotations from the Mouse ATAC atlas (displayed per rows) and the Cell-ID(g) cell type predictions (displayed per columns) using the gene-signatures extracted from the manually annotated cell types provided by the Tabula Muris mouse cell atlas. The color code in the heatmap represents the ratio *r* of the cell types displayed per rows that are allocated in the cell types represented per columns, ranging from white (*r* = 0) to red (*r* =1). (**D**) Global performance measured through the macro F1 score (y-axis) achieved by Cell-ID(g), Cell-ID(c) and 10 alternative state-of-the-art methods (x-axis) on the label transferring from the scRNA-seq to the scATAC-seq mouse cell atlas. DCT/CD: Distal convoluted tubule/ collecting duct, endo: endothelial, mono: monocytes.

Throughout this study, we found that Cell-ID was able to extract unbiased per-cell gene signatures from single-cell RNA-seq experiments that could then be used as unique cell identity cards without the need for prior knowledge. Cell-ID signatures were consistently reproducible, across diverse benchmarks collectively involving 14 independent single-cell datasets, 13 organs / tissues, more than 50 cell types, more than 200000 cells, 2 model organisms, 6 sequencing protocols and 2 single-cell-omics technologies. Such signatures made it possible to recognize cell identities across datasets from the same or different tissues of origin and model organisms, while overcoming batch effects arising from the use of different donors and sequencing technologies. In particular, the automatic cell-type annotations and matching across sets provided by Cell-ID were fully transparent with respect to the genes driving the hits. Such transparency improves biological interpretation at the individual cell level (**Supplementary Note 7**), making it possible to discover *bona fide* rare cells, as illustrated by the identification of Schwann cells and solitary chemosensory cells previously overlooked in published datasets (**Supplementary Note 3 and 4**). This contrasts strongly with the capabilities of the other methods currently available, which are based on assessments of similarity over the entire transcriptome^21^, cell embedding^22,29^, or machine-learning approach^23,24,26,27^, in which the contributions of individual genes are difficult to interpret.

Cell-ID is computationally efficient, can be scaled up for use with large datasets and allows *many-to-many* dataset comparisons without the need for data integration steps (**Supplementary Note 6**). It can, thus, be used for the systematic screening of each individual cell in newly sequenced datasets against (a) reference databases of marker lists associated with well-established cell types (e.g. PanglaoDB^37^, CellMarkers^38^), (b) reference single-cell atlas databases with manually curated cell-type annotations (Tabula Muris^35^, mouse ATAC atlas^36^, human cell atlas^1^), and (c) molecular signature collections, functional ontologies and pathway databases (e.g. MSigDB^39^, GO^40^, Reactome^41^, KEGG^42^, Wikipathways^43^). The automatic single-cell annotations provided by Cell-ID will alleviate the need for the often-tedious visual inspection of prototypic marker levels based on expert knowledge. From a discovery perspective, the identification of individual cells presenting distinctive and reproducible gene signatures consistent with the phenotypes studied would constitute a first step towards the in-depth experimental characterization of putative novel human cell types or cell states in health and disease. Cell-ID is implemented as an open-source R software package including detailed tutorials and scripts to reproduce all the analyses and figures presented in this manuscript (**Supplementary Software** and https://github.com/RausellLab/CelliD).

## Acknowledgements

We thank the Laboratory of Clinical Bioinformatics and the Laboratory of Human Lymphohematopoiesis for helpful discussions and support. The Laboratory of Clinical Bioinformatics was partly supported by the French National Research Agency (ANR) “Investissements d’Avenir” Program (Grant ANR-10-IAHU-01). The Laboratory of Clinical Bioinformatics and the Laboratory of Human Lymphohematopoiesis were partly supported by Christian Dior Couture, Dior. We also thank Dr. Gloria Fuentes from The Visual Thinker LLP for the creation of the illustrations in Figures 1-4.

## Author contributions

A.C. and A.R. conceived and designed research; A.C. performed research; A.C and L.M. contributed with materials/analysis tools; A.C and A.R. analyzed data; A.C, E.S and A.R. interpreted results; and A.C., E.S. and A.R. wrote the paper. All authors read and approved the final manuscript.

## Online methods

### Data availability

All single-cell data sets used in this paper are publicly available (**Supplementary Table 2**). scRNA-seq datasets for human blood cells profiled by Cite Seq^1^ and Reap Seq^2^ were downloaded from the gene expression omnibus (GEO; accession numbers GSE100866 and GSE100501, respectively). Cell-type labels for these two datasets were obtained following the Multimodal Analysis vignette of the *Seurat*^3^ R package (https://satijalab.org/seurat/multimodal_vignette.html). Pancreas scRNA-seq datasets from Baron^4^, Muraro^5^, and Segerstolpe^6^, as well as their associated cell-type annotations were downloaded via the scRNAseq^7^ R package as a SingleCellExperiment format R object. Plasschaert^8^ mouse and human and Montoro^9^ mouse airway epithelium scRNA-seq datasets, and their annotations were downloaded from GEO (GSE102580, GSE103354). Haber^10^ intestinal epithelium scRNA-seq dataset was downloaded from GEO accession code GSE92332. Olfactory epithelium scRNA-seq datasets from Fletcher^11^ and Wu^12^were downloaded from GEO (GSE95601, GSE120199), and their cell type annotations were obtained from the associated Github repositories: https://github.com/rufletch/p63-HBC-diff and https://www.stowers.org/research/publications/odr for Fletcher^11^ and Wu^12^, respectively. Tabula Muris^13^ 10X and Smart-seq mouse scRNAseq datasets were downloaded from https://tabula-muris.ds.czbiohub.org/. Gene activity score matrices from the Mouse sci-ATAC-seq atlas datasets from Cusanovich^14^ were obtained from http://atlas.gs.washington.edu/mouse-atac/data/, as provided by the authors and resulting from the aggregation of information across all differentially accessible chromatin sites linked to a target gene.

### Preprocessing and normalization of single-cell datasets

All single-cell RNA-seq datasets analyzed in the study took as input the raw count gene expression matrices provided by the original sources. Library size normalization was carried out by rescaling counts to a common library size of 10000. Log transformation was performed after adding a pseudo-count of 1. All analyses throughout the manuscript were restricted to a background set of 19308 and 21914 protein-coding genes from human and mouse, respectively, obtained from BioMart Ensembl release 100, version April 2020 (GrCH38.p13 for human, and GRCm38.p6 for mouse^15,16^). Genes expressing at least one count in less than 5 cells were removed. No filtering of cells was done unless the original sources provided “doublet” or contamination annotations, which were in such cased filtered out. In the case of sci-ATAC gene activity score matrices, the same preprocessing as for scRNA-seq data was applied.

### Overview of the Cell-ID approach

The main steps of the Cell-ID workflow are schematized in **Figure 1**. First, the library size normalized and log-transformed count matrix is transformed into a fuzzy coded indicator matrix where expression values are represented in a continuous scale between 0 and 1. Second, Cell-ID performs a dimensionality reduction of the indicator matrix using Multiple Correspondence Analysis where both cells and genes are represented into the same vector space^17^. Third, per-cell gene rankings are calculated from the gene-to-cell distances in MCA space, where the top closest genes to a cell will define its gene signature. If a grouping of cells is provided, per-group gene rankings may be obtained in an analogous way by using the geometric centroid in MCA space of the cells belonging to a given group. The enrichment of per-cell and/or per-group gene signatures is then evaluated through hypergeometric tests against (i) reference marker gene lists, and/or (ii) per-cell and/or per-group gene signatures extracted through Cell-ID from reference single-cell datasets. Per-cell and per-group gene signatures represent thus identity cards allowing automatic cell type and functional annotation, as well as cell matching across datasets. Each of these steps is described in detail in the following sections:

### Multiple Correspondence Analysis of the gene expression matrix

Multiple Correspondence Analysis (MCA) is a multivariate descriptive statistical technique conceptually equivalent to Principal Component Analysis for qualitative/binary data^18,19^. MCA can be applied to quantitative data through an intermediate step of a so-called *fuzzy* coding, Here, each continuous variable *p* is coded through user-defined functions into a number of disjoint categories *Q*_*p*_ where membership *x* to each category *q* is represented in a continuous scale between 0 and 1, and 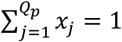. Following (Aşan and Greenacre, 2011), fuzzy-coding of a cases-by-variables matrix of continuous data can be performed in its simplest form by doubling each variable into *Q*_*p*_ = 2 categories as follows: Let *M*_*N,P*_ be the gene expression matrix of *N* cells (i.e. cases) and *P* genes (i.e. variables), with general term *m*_*np*_ gathering the expression level of gene *p* in cell *n*. For each column vector *Mp* in M, two membership functions *x*^+^ = *f*^+^(*m*): ℜ → ℜ: [0,1] and *x*^−^ = *f*^−^(*m*): ℜ → ℜ: [0,1], *where f*^−^(*m*) = 1 − *f*^+^(*m*), can be defined by linearly scaling between 0 and 1 the expression values for each gene across all cells as follows:

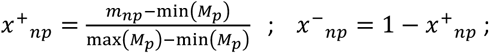

From such functions, a fuzzy-coded indicator matrix *X*_*N,K*_ can be built, representing a total of *K=2P* categories :

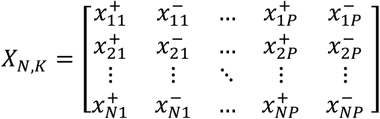

The grand total of *X*_*N,K*_ is thus *N*P*, since each of the *N* cells has P sets of fuzzy coded-values, each adding up to 1. The Multiple Correspondence Analysis (MCA) of a fuzzy-coded indicator matrix follows that of a regular MCA^20^. From the matrix *X*_*N,K*_, a matrix of relative frequencies *F*_*N,K*_ is defined as :

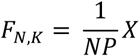

From the row sums and column sums of F, two diagonal matrices *D*_*r*_ and *D*_*c*_ are build, respectively, with general terms

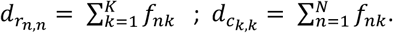

Let *S*_*NK*_ be the matrix of standardized relative frequencies resulting from

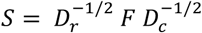

The singular-value decomposition (SVD) of the matrix *S*_*NK*_ leads to

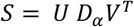

Where U and V contain by columns the singular vectors of norm 1 (U^T^U=1; V^T^V=1), and *D*_*α*_ is a diagnonal matrix of singular values ∝_*i*_, which are positive and displayed in descending order: ∝_1_≥ ∝_2_≥ … > 0. Alternatively, U and V can be obtained as the matrices displaying by columns the eigen vectors of norm 1 of the product SS^T^ and S^T^S, respectively, with eigen values *λ*_*i*_, where 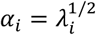. Thus,

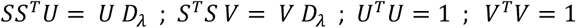

Alternatively, V can be calculated from U with the transition formula:

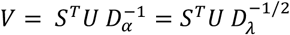

In the previous expressions, the first vectors 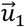 and 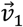 are associated with the trivial solution *α*_1_ = *λ*_1_ = 1; and are thus removed from the analysis at this stage. After eliminating the trivial solution, the sum of all the eigenvalues *λ*_*i*_ from SS’ (so-called t*otal Inertia* in MCA terminology) equals the Chi-squared statistic *χ*^2^ of the indicator matrix *X*_*N,K*_ divided by *N*. Thus, the orthogonal vectoral space generated by the eigenvectors of SS’ can be viewed as a decomposition of the Chi-squared statistic *χ*^2^ in its independent sources of variation, each accounting for a fraction given by 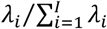. At this stage, further dimensionality reduction can be performed by retaining the first *J* eigen vectors as the most informative components, while disregarding the rest of dimensions from downstream analysis. Here we established *J* = 50 as the default parameter throughout all the analyses performed.

The orthogonal dimensions given by the eigenvectors U and V allow to simultaneous represent both rows (i.e. cells) and columns (i.e. gene categories) in *S* into the same orthogonal vectoral space, where coordinates are obtained as follows^21^:

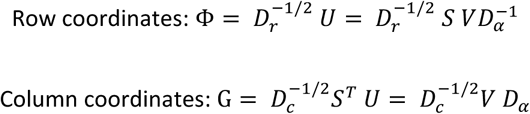

The previous expressions correspond to so-called *standard* and *principal* coordinates for rows and columns, respectively, in MCA terminology^21^. The simultaneous representation of cells and genes into the same vector space is a main advantage of MCA as compared to alternative dimensionality reduction techniques such as PCA, where only cells are projected.

### Gene-to-cell distances in MCA space and per-cell and per-group gene signature extraction

In the vectoral space provided by MCA, a barycentric relationship is fulfilled between the rows and the column coordinates: the general term *g*_*kj*_ of G representing the coordinate of a column *k* in the dimension represented by the eigenvector 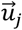 of U, corresponds to the weighted average (centroid) of the *N* row coordinates *ϕ*_*nj*_ from Φ, where weights are given by 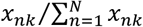, i.e. the frequency conditioned by columns of the corresponding values in the fuzzy indicator matrix *X*_*N,K*_ :

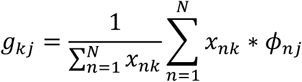

Thus, in MCA space the closer a column (i.e. gene category) is to a row (i.e. gene), the more specific it is to it. In addition, each set of *Q*_*p*_ = 2 categories for each gene *p* is centered at the origin: i.e. 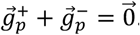. At this stage, only gene category coordinates 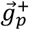, conveying presence of gene expression relative to the maximum per gene, are retained for downstream analysis. From the previous expressions, the Euclidean distances 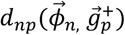 can be computed for each cell *n* and each gene *p* in the dataset. The genes *g*_*p*_ constituting the signature Γ_*n*_ associated to a cell *n* are obtained from its top *γ* closest genes in MCA space:

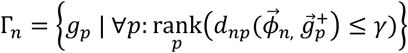

A default value of *γ* = 200 was established throughout this work and ties resolved with random ranks. More generally, the genes *g*_*p*_ constituting the signature Γ_Θ_ associated to a group of cells Θ can be obtained from the Euclidean distances 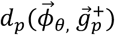 between each gene *p* and the group centroid 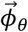, obtained from the geometric center of the 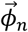 vectors associated to the cells *n* ∈ Θ.

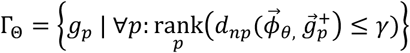

We note however that Cell-ID does not perform or relies in any clustering step whatsoever. Notwithstanding, cell grouping information may optionally be used here as input, as provided by a external reference source (e.g. database or publication).

### Per-cell gene signature enrichment analyses against reference gene sets

The gene signatures Γ_*n*_ extracted for each cell *n* in a dataset can be assessed through their enrichment against reference gene set (e.g. a marker gene list) associated to well characterized cell types and/or functional terms. Cell-ID evaluates such enrichment through a hypergeometric test as follows: Let *P* be the set of genes retained in the gene expression matrix *M*_*N,P*_ previously defined, after the initial steps of cell and gene filtering described above. Let *W* be the set of genes within a reference gene set which are contained on P (*W* ⊂ *P*). Let w be the number of genes overlapping between the signature Γ_*n*_ of size *γ* and the gene set *W*:

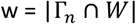

The observed overlap *w* can be modelled as a random variable *X* distributed hypergeometrically, with probability mass function given by:

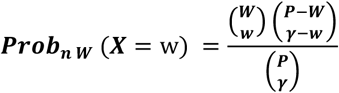

Only reference gene sets of size *W*≥ 10 were considered throughout this work. When the gene signature Γ_*n*_ of a cell *n* in a dataset *D* is evaluated against a collection of reference gene sets *W*_1_, *W*_2,_ … *W*_Ω_, (e.g. a repository of cell-type marker lists or a pathway database), the above hypergeometric test *p*-values are corrected by multiple testing for the number of gene sets Ω evaluated. Thus, a cell *n* is considered as enriched in those gene sets for which the hypergeometric test *p*-value is <1e-02, after Benjamini Hochberg multiple^22^ correction. In addition, when a disjointed classification is required, a cell *n* may be assigned to the gene set *W*_ω_ with the lowest significant corrected p-value. On the contrary, if no significant hits are found, a cell *n* will remain unassigned.

### Per-cell gene signature enrichment analyses against per-cell and per-group gene-signatures extracted from reference single-cell datasets

The gene signatures Γ_*n*_ extracted for each cell *n* in a dataset *D* can be assessed through their enrichment against the gene signatures Γ′_*n*_ extracted for each cell *n’* in a reference dataset *D’*, an approached called here Cell-ID(c). Analogous to the previous section, Cell-ID(c) evaluates such enrichment through a hypergeometric test as follows: Let *P* be the set of genes retained in the gene expression matrix *M*_*N,P*_ associated to dataset *D* as previously defined. Let Γ′_*n*′|*P*_ be the set of genes of size *W’* within a per-cell gene signature Γ′_*n*_ extracted for a cell *n’* in the dataset *D’*, which are contained on P, i.e.: Γ′_*n*′|*P*_ = Γ′_*n*′_∩*P*.

Let w be the number of genes overlapping between the signature Γ_*n*_ of size *γ* and the gene set P:

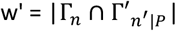

The observed overlap *w’* between two per-cell gene signatures can be modelled as a random variable *X* distributed hypergeometrically, with probability mass function given by:

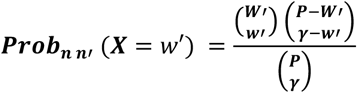

For each cell *n* in a dataset *D*, the above hypergeometric test *p*-values are corrected by multiple testing for the number of cells *N’* in the reference dataset D’ against which it is evaluated. Thus, a cell *n* in D is considered as enriched in those signatures n’ in D for which the hypergeometric test *p*-value is <1e-02, after Benjamini Hochberg correction on the number *n’* of tested gene signatures. In addition, when a disjointed classification is required, a cell *n* may be assigned to the cell n’ in D’ with the lowest significant corrected p-value. Best hits can be used for cell-to-cell matching and label transferring across datasets. On the contrary, if no significant hits are found, a cell *n* will remain unassigned.

Alternatively, if a grouping 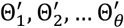 of the *N’* cells in D’ is provided, the gene signatures Γ_*n*_ for each cell *n* in a dataset *D* can be assessed through their enrichment against the corresponding per-group gene signatures 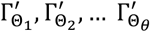 extracted from *D’* as described above. We call this approach Cell-ID(g). Here, a cell *n* in D is considered as enriched in those cell groups 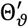 from D’ for which the hypergeometric test *p*-value is <1e-02, after Benjamini Hochberg correction for the number of groups evaluated. In addition, when a disjointed classification is required, a cell *n* may be assigned to the group 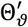 in D’ with the lowest significant corrected p-value. Best hits can be used for cell-to-group matching and group-based label transferring across datasets. On the contrary, if no significant hits are found, a cell *n* will remain unassigned. Cell-ID(g) can handle both disjoint and non-disjoint cell groupings (i.e. overlapping groups), as well as complete or non-complete groupings (i.e. when not all cells in D’ have been assigned to a group).

### Simulated datasets

Simulated scRNA-seq datasets were obtained with the Splatter^23^ Bioconductor package (version 1.4.1; https://bioconductor.org/packages/release/bioc/html/splatter.html). For the generation of structured datasets, each simulation was set to originate from five underlying subpopulations with relative sizes of 30%, 25%, 20%, 15% and 10% cells, respectively. No clustering or cluster labels were used in any way for the purpose of the analysis. Splatter’s logistic function, modeling the probability of a gene having zero counts, was defined by a midpoint parameter x0 = 3 counts, to obtain a simulated dataset with about 60% ∼ 70% of the count matrix content equal to zero after default normalization and filtering, a dropout rate consistent with those observed on other datasets used in this manuscript. Default values were used for all the other parameters. Centered and scaled principal component analysis (PCA) was performed with the base R prcomp function.

### Comparative benchmark of approaches performing marker-based cell type annotation

In the comparative benchmark assessing the method’s capacity to classify cells using pre-established gene signatures, two alternative semi-supervised classifiers were used: SCINA^24^ and AUCell^25^ (**Supplementary Table 3)**. Cell-ID predictions were based on a reference collection of blood cell markers from the XCell repository ^26^ (**Supplementary Table 4**). Only cell types of the hematopoietic lineage supported by more than three bulk RNAseq samples in XCell were considered: hematopoietic stem cells (HSC), multipotent progenitors (MPP), B cells, CD4+ T cells, CD8+ T cells, natural killer (NK) cells, plasmacytoid dendritic cells (pDC), common myeloid progenitors (CMP), granulocyte myeloid progenitors (GMP), megakaryocyte and erythrocyte progenitors (MEP), erythrocytes, megakaryocytes, platelets, basophils, eosinophils, neutrophils, CD14+ monocytes, CD16+ monocytes, macrophages, dendritic c(DC), conventional dendritic cells (cDC). For each cell type, the marker list used included genes replicated in at least 20% of the reported sources. Raw count matrices were used as input for AUCell and log-transformed normalized matrices were used for SCINA, following their associated vignettes. SCINA was run with default parameters except for i) the maximum number of iterations and the convergence rate, which were increased to 20 and 0.999 respectively to ensure a stable results, and ii) the rejection parameter, which was set to true to enable cells to be labelled as unassigned when there is a low confidence on the cell type prediction. For AUCell, the gene set with the highest AUC score was used to classify cells unless the maximum AUC score was <0.1, what left a cell as ‘unassigned’.

### Comparative benchmark of approaches performing cell-type label-transferring across datasets

In addition to Cell-ID, we evaluated 10 alternative approaches for cell-type label-transferring across scRNA-seq (**Supplementary Table 3**). All methods were run using default parameters unless otherwise stated. When default settings were not explicitly defined, setting used in the associated vignettes were followed. Methods used as input either the raw or the normalized count data (after gene filtering as described above), following each method’s documentation. For those methods that stipulate it, gene expression matrices were further restricted to genes in common between the reference and the query datasets. In the case of MNN^27^, we transferred labels from the reference to the query datasets between closest mutual nearest neighbor cells, and cells were left unassigned when no mutual nearest neighbor cells were found. We modified the default parameter of k nearest neighbor of MNN to k = 50 as the default k = 20 failed to find mutual nearest neighbor matches for a large fraction of cells, which negatively affected the benchmark metrics evaluated. Furthermore, for the selection of hypervariable genes in MNN, we used the default Seurat function for highly variable gene detection (2000 genes), and we took the intersection between the reference and the query highly variable genes to perform the query and reference dataset integration following the package’s vignette. In the case of SCN^28^, all cells that were classified as *rand* were considered as unassigned, as well as all cells classified as “nodes” in CHETAH^29^. For Seurat cells were labelled as unassigned when the projection score was below 0.5.

### Classification performance assessment

Cell annotation performance was assessed through 3 complementary metrics i.e.: precision, recall and F1 score, which is the harmonic mean between the precision and recall. Each metric was first calculated for each cell type in the query dataset. Second, each metric (i.e. recall, precision and F1 score) was calculated for the global set as the arithmetic mean of the corresponding metric across the evaluated cell types. In such a way, an overweighed contribution of largely populated cell types is avoided, allowing thus a larger contribution of more rare cell types to the metrics evaluated. Mapping across datasets of cell type nomenclature was performed through manual curation and is reported in **Supplementary Table 5**. For label-transferring performance assessment, cells left unassigned in a query dataset were considered as a false assignment when their actual cell type was indeed represented in the reference dataset, or considered as a true assignment otherwise. Therefore, for intestinal dataset label transferring, any cell types apart from endocrine, brush/tuft and goblet were considered as negative cell type since there are not present in the reference dataset and hence should be labeled unassigned. Thus, unassigned cells were considered as true positive if labelled against negative cells and the negative cells were evaluated in the calculation of performance metrics in the same manner as the three other cell types present in the reference and was considered also in the global performance metrics. For interspecies label-transferring across human and mouse datasets, the initial raw matrix of the query dataset was restricted to genes with one-to-one orthologs. Ortholog relations were obtained from BioMart (release 100, version April 2020, GrCH38.p13 for human, and GRCm38.p6 for mouse^15,16^) using gene symbols.

### Functional Enrichment

Functional enrichment analyses were performed using 6 sources of functional annotations: Reactome^30^, KEGG^31^, WikiPathways^32^, GO biological process^33^, GO molecular function and GO cellular component, collectively gathering a total of 8709 terms. Gene sets associated to functional pathways and ontology terms were obtained as provided in enrichr^34^ website http://amp.pharm.mssm.edu/Enrichr/#stats. (**Supplementary Table 6**)

### Visualization

UMAP^35^ or tSNE^36^ representations were alternatively used for visualization purposes throughout the manuscript. However, no biological conclusions whatsoever were drawn from visual inspection of such representations. UMAP and tSNE representations were obtained with Seurat default parameters following this vignette (https://satijalab.org/seurat/v3.1/pbmc3k_tutorial.html).

### Computational resources

All analyses presented in the manuscript were run in a workstation with 64-GB RAM memory and a AMD Ryzen 2700X processor with 8 3.6-GHz physical cores, with the exception of the scalability benchmark presented in **Supplementary Note 6** where an Intel Xeon Gold 6140 with 36 2.3 GHz cores processor and 640Gb of RAM was used.

### Code availability

Cell-ID is implemented as an R package and is available on GitHub (https://github.com/RausellLab/CelliD) under the GPL-3 open source license. Complete documentation is provided with step-by-step procedures for MCA dimensionality reduction, per-cell gene signature extraction, cell type prediction, label-transferring across datasets and functional enrichment analysis. All R scripts and intermediate data representations required to reproduce all figures in the manuscript are provided as a supplementary file.

## Supplementary Notes 1 - 7

**Supplementary Note 1. Consistency of MCA low-dimensional representation of cells and genes**.

The consistency of MCA-based low-dimensional representation of cells and genes was first evaluated with 100 simulated scRNA-seq datasets, each containing 1000 cells and 5000 genes (**Online Methods**). We first compared the correspondence between MCA and PCA low-dimensional representations of cells by performing Spearman’s rank correlation analysis on their principal axes coordinates. MCA and PCA cell representations were largely equivalent (**Supplementary Fig. 1**). We then determined whether the per-cell gene rankings obtained with MCA were consistent with the gene expression values for neighboring cells in the MCA space. As expected, genes specific to a given cell had higher log-fold changes in expression in the 5% of cells closest to target cell (*n* = 50) than in the other cells (**Supplementary Fig. 2A)**. We then investigated how the ranking of genes with zero-counts in a cell related to the specificity of the genes concerned in neighboring cells (**Supplementary Fig. 2A)**. We found that genes not detected in a given cell were nevertheless attributed a high ranking in this cell if the surrounding cells displayed high levels of expression for these genes (**Supplementary Fig. 2A)**. This is an important result, highlighting the capacity of multivariate approaches to consider a gene to be specific to a cell in which it was not detected, provided that the gene concerned is specific to very similar cells. The MCA approach is thus robust to zero-count values that probably correspond to technical dropouts. These results could be generalized to all individual cells in a given dataset: Spearman’s rank correlation coefficient between rank and log-fold change = 0.72, *p*-value <10e-16 (**Supplementary Fig. 2B)**. By contrast, the correlation was weaker for per-cell gene rankings obtained via a naïve approach based on either (i) the log-fold changes in gene expression observed in one cell relative to all other cells (Spearman’s rank correlation coefficient = 0.38, *p*-value <10e-16, **Supplementary Fig. 2, C-D)** or (ii) highest-to-lowest rankings of expression values within a cell, with random ranks for ties, as previously described (AUCell^1^, Spearman’s rank correlation coefficient = 0.08, *p*-value = 3.18e-04, **Supplementary Fig. 2, E-F**, respectively).

**Supplementary Note 2. Capacity of Cell-ID to identify well-established cell types on the basis of reference marker lists**.

We evaluated the ability of Cell-ID per-cell gene signatures to reproduce reference marker lists associated with well-established cell types. We searched for scRNA-seq datasets in which concomitant measurements of single-cell protein marker levels had been performed. Protein marker levels provide additional evidence for cell-type annotations. Cell-type labels in such settings can be taken as a gold standard reference for this study, thereby avoiding the potential circularity associated with the use of transcription-based annotations. Two independent scRNA-seq datasets for human blood cells profiled by CITE-seq^2^ and REAP-seq^3^ protocols met this criterion (**Online Methods**). The Cite-Seq dataset contained data for a total of 8005 human cord blood cells whereas the REAP-seq dataset contained data for 7488 PBMCs. Cell-ID predictions were based on a reference collection of blood cell markers from the XCell^4^ repository (**Supplementary Table 4, Online Methods**).

The gene signature of each cell, extracted from the CITE-seq and REAP-seq datasets, was first evaluated with Cell-ID against each of the 21 marker lists associated with previously described cell types (**Figure 2** and **Supplementary Figure 3, Supplementary Table 4**; **Online Methods)**. Each cell was assigned to the cell type for which it displayed the strongest enrichment (**Online Methods**). We assessed the accuracy of Cell-ID by calculating its precision (positive predictive value), recall (true positive rate), and overall agreement (F1) with the previously assigned reference cell-type labels. In both datasets, per-cell gene signatures displayed significant enrichment in the marker genes for at least one reference cell type in 83% and 73% of cells for CITE-seq and REAP-seq datasets, respectively. The best match to the gene signature was used for automatic cell-type prediction with high precision (0.87 and 0.9), recall (0.84 and 0.73), and F1 (0.83 and 0.78) values (multinomial *p*-value < 10^−16^ for all figures; **Figure 2, Supplementary Figure 4** and **Supplementary Table 7**). Cell-ID outperformed reference methods for cell-type classification based on pre-established signatures, such as SCINA^5^ and AUCell^1^, for most of the cell types considered, achieving remarkably high levels of precision. CellAssign^6^ could not be used due to execution errors of the original software, which we attribute to the length of the signatures used here, ranging from 13 to 174 genes in total, which is much longer than the gene sets of 2 to 20 genes generally used by CellAssign. Overall performance was better for the CITE-seq dataset than for the REAP-seq dataset, for all methods evaluated, due to a significant difference in the median number of genes detected per cell between the two datasets (2346 and 1260, respectively, two-sided Wilcoxon test *p* value < 10^−16^).

In addition to identifying the best match between gene lists, Cell-ID was able to identify cells displaying significant enrichment in the genes of more than one of the pre-established cell-type marker lists. This contrasts with clustering-based approaches, which partition cells into disjointed groups. This situation is illustrated by the CITE-seq CBMC dataset, for which Cell-ID identified fine-grained transitions within the Hematopoietic Stem and Progenitor cells subset (CD34^+^) according to the hematopoietic hierarchy **(Figure 2C)**. No immature CD34^+^ cells in the REAP-seq dataset were reported in the original publication ^3^. The gradient of Cell-ID enrichment scores was consistent with the Hematopoietic Stem Cells (HSC) lineage differentiation process. Thus, Cell-ID multi-class scores reflect smooth transitions at individual cell level from the most immature HSC to multipotent progenitors (MPP), branching between myeloid (CMP/GMP) and erythroid (MEP) progenitors, which ultimately differentiate into erythrocytes. Such fine-grained transitions would have been missed by both clustering-based approaches and alternative methods. Thus, SCINA and AUCELL provided a coarse-grained classification between HSCs and erythrocytes, with no assignment to intermediate states between these two classes.

We then analyzed more challenging scenarios based on real datasets in which we evaluated the capacity of Cell-ID to identify the only cell of a given rare cell type (*n*=1). We focused on cell types observed at low frequencies (<2%) in the previous CBMC dataset: plasmacytoid dendritic cells (pDC, 0.6%, *n*=49), erythrocytes (1.3%, *n*=105) and hematopoietic stem and progenitor cells (HSPC, CD34+ subset, 1.8%, *n*=134) from the CITE-Seq dataset. For each of these rare cell types containing a given number *n* of cells, we generated *n* corresponding datasets, each retaining only one cell at a time, while the rest of the subpopulations remained unchanged. Using the pre-established immune cells marker lists, as described above, Cell-ID identified the singleton cell with a high mean recall (92%, 94%, and 86%), but modest precision (32%, 8% and 76%) due to the experimental design with very high number of negatives compared to positive (1 positive and 999 negatives), which results in mid-range F1 score (48%, 0.15%, 0.81%) (**Supplementary Table 8**). Singleton cells pose a problem in clustering-based approaches, in which they are often merged with larger sub-populations or treated as outliers and are thus filtered out of downstream analyses. Conversely, alternative reference methods able to work at individual cell level without a clustering step, such as SCINA^5^ and AUCell^1^, could not be applied in this setting either, because they are not suitable for independent evaluations one cell type at a time.

**Supplementary Note 3. Capacity of Cell-ID to match cells of analogous cell types across independent scRNA-seq datasets from the same tissue of origin, within and across species**.

Cell-ID provides two options for cell type matching across datasets: (i) Cell-ID(c), in which individual cells in a *query* dataset are matched to transcriptionally analogous cells in a *reference* dataset; and (ii) Cell-ID(g), in which individual cells in a *query* dataset are matched to cell group labels previously established in a *reference* dataset through clustering and/or expert annotation. Thus Cell-ID(c) provides *cell-to-cell* matching, whereas Cell-ID(g) performs *group-to-cell* matching from *reference-to-query datasets*. It should be noted, however, that Cell-ID(g) performs no clustering whatsoever, as the cell groups in the *reference* dataset are provided as an input by the *reference* source. For the evaluation of Cell-ID(c) and Cell-ID(g), we searched for tissues profiled in several independent scRNA-seq datasets and meeting the following three conditions: (i) cell type labels curated through expert annotation in the original publications, (ii) containing at least one rare cell type (cell types accounting for less than 2% of the cells in a sample), and (iii) different sequencing protocols and/or different model organisms (i.e. humans and mice) used. Two tissues meeting these requirements were identified (**Figure 3**):

(A) Pancreatic islets: we considered three scRNA-seq datasets for pancreatic cells from human donors: (i) 8659 cells from four deceased donors healthy at the time of death profiled with the inDrop protocol (Baron et al.^7^), (ii) 2126 cells from four deceased organ donors sequenced with CEL-seq2 (Muraro et al.^8^) and (iii) 2168 cells from six healthy donors and four donors with type-2 diabetes sequenced with Smart-seq2 (Segerstolpe et al.^9^). The Baron dataset reported the greatest diversity of cell types and was, thus, chosen as the *reference* for this analysis. The cell types identified in Baron’s human dataset included two exocrine cell types (acinar and duct cells), five endocrine cell types (alpha, beta, delta, gamma and epsilon cells), three immune cell types (tissue-resident macrophages, mast cells, cytotoxic T cells), pancreatic stellate cells, endothelial cells and Schwann cells of neural crest origin. All of these cell types, with the exception of Schwann cells and cytotoxic T cells were reported by Muraro et al. and Segerstolpe et al.

(B) Airway epithelium: three independent airway epithelial cell datasets from mice and human donors were obtained, including (i) 7662 tracheal epithelial cells from four mice profiled with 10X Genomics technology (Plasschaert et al^10^); (ii) 7586 airway tracheal cells from six mice sequenced with inDrop (Montoro et al.^11^); (iii) 2970 primary bronchial epithelial cells from three human donors profiled with 10X Genomics technology (Plasschaert et al^10^) (**Online Methods**). All three datasets reported the six major airway epithelial cell types, including basal, secretory cells (also known as club cells), ciliated cells, rare pulmonary neuroendocrine cells (PNEC), brush cells (also known as tuft cells) and ionocytes. Plasschaert’s mouse dataset was used as the *reference* for this study, as it allowed a within-species analysis (mouse-to-mouse) as well as a between-species analysis (mouse-to-human) based on datasets originating from the same laboratory.

Cell-ID gene signatures at either individual cell level or group level were obtained independently for each of the datasets described above (**Online Methods**). Four independent *reference*-to-*query* assignments were evaluated: Baron’s human pancreatic cells against Muraro’s and against Segerstolpe’s human pancreatic datasets, and Plasschaert’s mouse airway epithelial cells against Montoro’s mouse airway epithelial cells and against Plasschaert’s human airway epithelial cells. For each of the *reference*-to-*query* assessments, both *cell-to-cell* matching, and *group-to-cell* matching were performed, with Cell-ID(c) and Cell-ID(g), respectively (**Methods**). Thus, each cell in the query dataset was assigned to the cell type from the *reference* cell or *reference* group for which it presented the lowest significant gene signature enrichment *p* value (or was left unassigned if no significant hits were found; **Supplementary Figure 5**). Cell-ID *cell-to-cell* matching and *group-to-cell* matching were thus evaluated by assessing the cell-type agreement between the transferred labels and the original labels in the *query* dataset. Comparative benchmarking was performed against a representative set of alternative state-of-the-art methods covering major approaches for cell-matching or label transfer across scRNA-seq datasets (**Supplementary Table 3**): (i) integration-based methods for cell-to-cell matching across datasets based on reciprocal *k*-nearest neighbor analysis and projection in a common low-dimensional space (Seurat^12^, MNN^13^); (ii) transcriptome similarity assessment without an integration step (scmap_cluster and scmap cell^14^, scID^15^ and singleR^16^); and (iii) machine learning-based methods training models through cross-validation in the reference dataset followed by prediction in the query dataset (CaSTle^17^, scPred, SCN^18^and CHETAH^19^).

Both Cell-ID(c) and Cell-ID(g) consistently reached high precision (>82% and >83%), recall (>82% and >=76%) and F1 values (>74% and >74%, respectively) across all *reference*-to-*query* assignments evaluated (multinomial *p* value < **2.2e-16** for all figures; **Figure 3A, Supplementary Figure 6 A-B, Supplementary Table 9**). Cell-ID performance was at least as good as that of all alternative state-of-the-art methods. Notably, Cell-ID obtained high performance scores even in for cell-type matching across species, which proved challenging for most of the methods evaluated. We assessed the robustness of Cell-ID gene signatures further by focusing on the cell types present at low frequencies (<2%) in the previous query datasets: epsilon cells, tissue-resident macrophages, mast cells and endothelial cells from pancreatic islet samples (Muraro et al. and Segerstolpe et al), and PNEC, brush cells and ionocytes in the mouse and human airway epithelium datasets (Montoro et al and Plasschaert et al., respectively). As expected, *reference-to-query* assignments for such rare cell types were generally more error-prone for all the methods evaluated, including Cell-ID (**Figure 3B, Supplementary Figure 6 C-D**). Nevertheless, Cell-ID performed better than the alternative methods, with salient scores for Cell-ID(g) for most of the rare cell types evaluated (median F1 values greater than 88%, respectively, with *p* value<1e-5 for all evaluated rare cell populations relative to expectations from a random binomial distribution).

From a discovery perspective, the datasets used here provided us with an opportunity to evaluate the capacity of Cell-ID to detect ultra-rare cell types potentially missed in the original publications. Indeed, Baron et al. reported the presence of Schwann cells (*n*=13; 0.17 %) in pancreatic islet datasets, an ultra-rare cell type of neural crest origin. However, Schwann cells were not described in the samples of Muraro et al. and Segerstolpe et al. We thus explored possible enrichment in the Cell-ID gene signatures extracted for each of the 13 Schwann cells in the Baron et al. dataset for the 2126 and 2168 cells, respectively of the query datasets. Both Cell-ID(c) and Cell-ID(g) identified *n*=4 cells in the Muraro et al. dataset and n=2 cells in the Segerstolpe et al. dataset that had been initially labeled as ductal (Muraro et al.^8^) or unclassified (Segerstolpe et al.^9^) in the associated publications. These cells presented high expression levels of Schwann cell markers (SOX10, S100B, CRYAB, NGFR, CDH19, and PMP22) and of markers of response to nerve injury (SOX2, ID4, and FOXD3), as described by Baron et al.^7^. Accordingly, SOX10, S100B, CRYAB, NGFR, CDH19, and PMP22 ranked among the top 100 gene signatures associated with the four and two cells annotated as Schwann cells by Cell-ID (**Supplementary Figure 7**), and these six genes were significantly upregulated in these cells relative to the other cells (Wilcoxon test *p* value < 10^−3^, in both datasets). The identification of putative Schwann cells was replicated by Seurat, MNN, scID, SCN, SingleR and scmap cluster, but missed by CaSTLe, scmap cell, scPred and CHETAH, which labeled them as “other” or “unassigned”. Moreover, functional enrichment analysis of the gene signatures of each of the four and two putative Schwann cells revealed significant enrichment in 62 Gene Ontology (GO) biological processes (median p-value across cells <0.01) for the Muraro et al. dataset and 26 GO biological processes for the Segerstolpe et al. dataset, with 10 terms common to the two sets (**Supplementary Table 1**). Nervous system development (GO:0007399), including glial cell differentiation (GO:0010001) and axonogenesis (GO:0007409), were among the top 10 terms displaying enrichment in both sets (p-values less than or equal to 2.18e-05 and 2.98e-03, respectively), consistent with the neural crest origin of Schwann cells^20^. The prominent role played by Schwann cells in the myelination process ^21^ was, in turned, reflected in several functional terms and pathways for which significant enrichment was detected, including collagen fibril organization (GO:0030199) in the four cells from the Muraro dataset and myelination (GO:0042552) and collagen binding (GO:0005518) in the two cells from the Segerstolpe dataset. These signals were consistently reproduced with alternative pathway annotation sources such as KEGG^22^, Reactome^23^ and WikiPathways^24^ (**Supplementary Table 1, Supplementary Table 6**).

Overall, these results show that Cell-ID gene signatures are highly reproducible at both the cell and group levels, across independent datasets from the same tissue of origin, despite the use of different sequencing protocols, donors, and species. This reproducibility was robust even for rare cell types present at extremely low frequencies, demonstrating the ability of Cell-ID to extract biologically relevant gene signatures at individual cell resolution. In particular, matching across datasets by Cell-ID is fully transparent in terms of the set of genes driving the hits, and is therefore fully interpretable in biological terms. This contrasts sharply with the situation for label transfer methods based on assessments of similarity over the entire transcriptome (e.g. Seurat, MNN, scmap) or machine-learning approaches, for which individual gene contributions are difficult to interpret (e.g. scPred, SCN).

**Supplementary Note 4. Capacity of Cell-ID to match cell types across independent scRNA-seq datasets from different tissues of origin**

In a more challenging scenario, we then evaluated the capacity of Cell-ID to recognize gene signatures of rare cells across independent sets from different tissues of origin. We thus searched for rare cell types meeting the following conditions: (i) that they were common to at least two different tissues, and (ii) for which single-cell RNA-seq datasets were available, in which cell-type labels had been curated by expert annotation in the original publications. Chemosensory epithelial cells appeared to be an ideal rare cell type present in various epithelium types, presenting all the characteristics for the purpose of the analysis. Recent studies have shown that chemosensory epithelial cells, referred to as tuft cells in the intestinal mucosa, and solitary chemosensory cells (SCCs) in the nasal respiratory mucosa, belong to the brush/tuft cell family, together with the brush cells in the tracheal airway epithelium ^25 26^. This chemosensory epithelial cell family has common transcriptional programs and functions in all three tissues ^27^. In particular, these cells are the primary source of interleukin-25^28^, a proinflammatory protein mediating type 2 inflammation induced by diverse pathogens in various mucosal tissues.

We evaluated the capacity of Cell-ID to identify rare cell types across tissues, by focusing on cell matching between the mouse tracheal airway epithelium (Plasschaert ^10^ and Montoro ^11^ datasets, as described in **Supplementary Note 3**) and the mouse small intestinal epithelium (single-cell dataset from Haber et al^29^, for 7216 cells from four mice sequenced with the inDrop protocol). The mouse airway and intestinal epithelia display major differences in terms of their cell-type composition. Both Plasschaert et al. and Montoro et al. reported the presence of basal, secretory (including goblet in Montoro), ciliated cells, PNEC, ionocytes and brush/tuft cells in the airway epithelium (**Supplementary Note 3**). Haber et al. performed single-cell analyses on the intestinal epithelium in which they identified enterocytes (45%), transit-amplifying cells (21%), stem cells (18%), goblet cells (7%), Paneth cells (4%) enteroendocrine cells (4%) and brush/tuft cells (2%) (**Supplementary Figure 8A)**. Cell-ID(g) and Cell-ID(c) were applied with default parameters (**Online Methods)** for *reference*-to-*query* assignments from Plasschaert and Montoro’s mouse airway epithelial cells to Haber’s mouse intestinal epithelial cells. Each cell in the mouse intestinal epithelial dataset was assigned to the cell type from the mouse airway epithelial for which it presented the lowest significant *p* value for gene signature enrichment (or was left unassigned if no significant hits were found). Both Cell-ID(g) and Cell-ID(c) identified brush/tuft, but also endocrine cells and goblet cells in the intestinal epithelium, based on brush/tuft, PNEC and secretory/goblet gene signatures, respectively, extracted from the airway epithelium (**Figure 3C** and **Supplementary Figure 8B**). Global matching accuracy was evaluated by analyzing the concordance of cell-type labeling with the original cell-type annotations from the corresponding publications (**Supplementary Table 5**). Comparative benchmarking was performed against a representative set of alternative state-of-the-art methods covering major approaches for label transfer or cell-matching across scRNA-seq datasets (**Supplementary Table 3**), as previously described in **Supplementary Note 3**. Cell-ID(g) yielded high precision, recall and F1 scores, clearly outperforming all the other methods evaluated (**Figure 3D, Supplementary Figure 8C** and **Supplementary Table 10)**. Similar results were obtained when the Plasschaert or Montoro mouse airway epithelial cells were used as the *reference set*. Alternative methods, including Cell-ID(c), performed less well, by contrast to the results obtained for label transfer between samples from the same tissue of origin (**Supplementary Note 3**). This lower performance was driven by many false positives (i.e. incorrect label transfer from the reference to the query dataset), resulting in low precision and low F1 scores. By contrast, correct label assignment rates remained high for Cell-ID(g), which did not assign labels from the reference to the query set in the absence of significant statistical support. The higher performance of Cell-ID(g) than of Cell-ID(c) suggests that group-based gene signatures are more robust than individual cell-based signatures for cell-type matching across samples from different tissues of origin.

We then searched for olfactory epithelium single-cell datasets of use for the evaluation of the capacity of Cell-ID to identify solitary chemosensory cells (SCCs) by projecting brush/tuft signatures extracted from the airway and intestinal epithelia. Two datasets from published works were identified: (i) Wu et al^30^, in which a total of 9126 olfactory epithelium cells from mice aged 0, 3, 7 and 21 days were sequenced with the 10X Genomics protocol, and (ii) Fletcher et al.^31^, in which 849 cells from mice aged 21-28 days were sequenced with the Smart-Seq2 protocol. The mouse olfactory epithelium has a cell-type composition different from those of the airway and intestinal epithelia. Both Wu et al and Fletcher et al reported the presence of transitional horizontal basal cells, resting horizontal basal cells, microvillous cells, mature sustentacular cells, mature olfactory sensory neurons, immediate neuronal precursors, immature sustentacular cells, immature olfactory sensory neurons, globose basal cells and unknown/unlabeled cells. However, neither of these studies reported the presence of SCCs among the cells sequenced. Nevertheless, these two datasets provided us with an opportunity to illustrate the capacity of Cell-ID for exploratory cell-type scanning on single-cell datasets of a *query* tissue, using gene signatures extracted from single-cell datasets for a set of *reference* tissues. Cell-ID(g) was used to evaluate the presence in the olfactory epithelium of cells with the tuft/brush cell gene signatures extracted from the **airway** and **intestinal epithelium**, as described above. Cell-ID(g) identified a total of 37 cells in the Wu dataset and 5 cells in the Fletcher dataset displaying gene signatures with significant enrichment in the genes of the tuft/brush cell signatures extracted from the **airway epithelium** (dataset from Plasschaert et al^10^). Cell-ID(g) also identified the same 37 and 5 cells as having signatures significantly enriched in the genes of the tuft/brush cell signatures extracted from the intestinal **epithelium** (dataset from Haber et al). Interestingly, these cells were labeled “unidentified” in the original publications ^30 31^. They displayed higher levels of expression for IL25 and the chemosensory cell marker Gnat3 than any of the other cells in the corresponding datasets (two-tailed Wilcoxon test, *p* <10-16 for both datasets), further supporting the classification of these cells as SCCs (Ualiyeva et al. 2020; **Figure 3E-F, Supplementary Figure 9)**. Moreover, functional enrichment analysis performed on the gene signatures of each of the 37 and 5 putative SCCs identified 20 and 14 Gene Ontology (GO) biological processes (median *p* value across cells <0.01) displaying significant enrichment in the datasets of Wu and Fletcher, respectively, with nine terms common to the two datasets (**Supplementary Table 1**). Unsaturated fatty acid biosynthetic process (GO:0006636), including eicosanoid (GO:0046456) and leukotriene (GO:0019370) biosynthetic processes, were among the top five terms displaying the highest level of enrichment in both datasets (5.76E-04 and 1,77E-05, respectively), driven by the Alox5, Alox5ap, Ltc4s, Pla2g4a genes in both sets of putative SCCs. This finding is consistent with recent studies showing that SCCs are a primary source of cysteinyl leukotriene in the olfactory epithelium ^25 26^), and that these cells have immune effector functions in common with airway brush/tuft cells ^25^. These signals were consistently reproduced with alternative pathway annotation sources (KEGG^22^, Reactome^23^ and WikiPathways^24^; **Supplementary Table 1**). Taken together, the gene signatures and functional hallmarks revealed by Cell-ID(g) confirm, at single-cell transcriptional level, the presence of rare cells in the olfactory epithelium similar to the airway and intestinal brush/tuft cells. In accordance with recent findings ^25 26^, the 37 cells from Wu et al and the five cells from Fletcher et al described above should, thus, be annotated as SCCs.

**Supplementary Note 5. Capacity of Cell-ID to match cell types across independent datasets from different single-cell *omics* technologies: scRNAseq and scATACseq**

We investigated whether the gene signatures extracted with Cell-ID from scRNA-seq datasets were replicated in independent single-cell ATAC-seq experiments on the same or different tissues of origin. We made use of two recent large-scale single-cell projects annotating cell-type heterogeneity through in-depth expert curation in a comprehensive set of organs and tissues for the model organism *Mus musculus*: the Tabula Muris^32^ mouse cell atlas, based on scRNA-seq, and the Mouse ATAC Atlas^33^, based on scATAC-seq. The Tabula Muris project profiled 20 mouse organs and tissues in a total of eight adult mice, with two alternative sequencing technologies: SMART-Seq2 (53760 cells) and 10X genomics (55656 cells). The Mouse ATAC Atlas reported genome-wide chromatin accessibility in ∼100,000 single cells from 17 samples spanning 13 different tissues in 13 adult mice. Only cells from the eight organs/tissues common to the Tabula Muris and ATAC Atlas datasets were retained for downstream analyses: heart, kidney, liver, lung, bonne marrow, spleen, thymus and large intestine (large intestine available only for SmartSeq scRNA-seq and scATACseq). Collectively, the tissues retained contained 50, 43 and 37 different cell types for the 10X Genomics, SMART-Seq2 and scATAC-seq datasets, respectively. Cell-type nomenclature equivalences between the Tabula Muris and the ATAC Atlas datasets are presented in **Supplementary Table 5**. Comparisons were made across datasets based on gene expression matrices (scRNA-seq) and gene activity scores (scATACseq) (**Online Methods**).

Cell-ID(g) and Cell-ID(c) were applied with default parameters (**Online Methods)** for *reference*-to-*query* assignments from (i) SMART-Seq2 and (ii) 10X genomics Tabula Muris scRNA-seq data to the Mouse ATAC Atlas. Thus, each cell in the Mouse ATAC Atlas was assigned to the cell type from the Tabula Muris scRNA-seq dataset for which it presented the lowest significant *p*-value for gene signature enrichment (or was left unassigned if no significant hits were found). We performed this procedure independently for the SMART-Seq2 and 10X genomics datasets. Global matching accuracy was evaluated by assessing the concordance of cell-type labeling between the manually curated cell type annotations provided by the corresponding databases (**Supplementary Table 5**). Comparative benchmarking was performed against a representative set of alternative state-of-the-art methods, as described in **Supplementary Note 3 and 4**, except for scID^15^, which could not be included here due to computational time limitations (>48 h in our computing infrastructure; **Methods**). By detecting shared gene signatures, both Cell-ID(g) and Cell-ID(c) correctly matched equivalent cell types across scRNA-seq and scATAC-seq datasets (**Figure 4, Supplementary Figure 10-12)**. Thus, both Cell-ID(g) and Cell-ID(c) maintained a good balance between precision and recall rates, resulting in high F1 scores (>0.74 for 10X and >0.6 for SmartSeq2), clearly outperforming all other methods evaluated, with the exception of SingleR, which yielded slightly lower performances (**Supplementary Figure 12, Supplementary Table 11)**. Overall, these results confirm that Cell-ID can extract, in an unbiased manner, gene signatures that are robustly reproduced across diverse single-cell *omics* technologies applied across highly heterogeneous cell types from multiple tissues and organs.

**Supplementary Note 6. Computational details, timing and memory consumption**

To evaluate the Cell-ID scalability to massive single-cell RNA-seq dataset analysis, we evaluated its time and memory consumption for different input sizes, and benchmarked them against the state-of-the-art methods previously considered in our study (**Supplementary Notes 3, 4** and **5)**. To that aim, we performed large-scale cell mapping and label transfer tasks between different *reference-to-query* datasets using single-cell RNA-seq data from the Tabula Muris atlas (see **Supplementary Note 5**). All 20 tissues and organs were considered here. For the purpose of the analysis, we first randomly subset an increasing number of cells (200, 500, 1000, 2000, 5000, 10000, 20000 and 50000 cells) from the 10X and -independently-from the Smart-seq datasets. Sampling of the reference dataset was restricted to the 10 most abundant cell types from 10X. We fixed the dataset to have 10 different subpopulations since the number subpopulations in the reference dataset impacts significantly computation time for some methods such as scID or SingleR. We then evaluated the computation time and the total memory allocation of a label transfer task as a function of the number of cells in the *query* dataset. Thus, label transfer from a fixed *reference* subset of 5000 cells from the SmartSeq dataset was performed against 8 different *query* datasets of increasing size, corresponding to the 8 subsets from the 10X Genomics dataset previously described (**Supplementary Figure 13 A-B**). In an analogous manner, we evaluated the computation time and the total memory allocation of a label transfer tasks as a function of the number of cells in the *reference* dataset. Here, label transfer from 8 different *reference* datasets of increasing size (corresponding to the 8 subsets from the Smart-seq dataset previously described) against a fixed *query* subset of 5000 cells from the 10X Genomics dataset (**Supplementary Figure 15 C-D**). Overall, Cell-ID computational time behaved in a comparable way to other state-of-the-art methods, yet with slightly higher memory consumption. The previous benchmark is based on the cell matching and label transfer between a pair of datasets. In the case of Cell-ID, such process involves (i) the MCA low dimensionality reduction and per-cell gene signature extraction for each dataset, and (ii) cell-to-cell matching between datasets, based on hypergeometric tests on the pre-extracted per-cell gene signatures. From a computational point of view, each of such cell-to-cell evaluations constitutes a fully independent job, and thus they can be readily parallelized in a multi-core and multi-node computing cluster. Cell-ID thus allows the creation of reference libraries of individual cell’s gene signatures from massive collections of single-cell RNA-seq datasets, enabling efficient large-scale cell-to-cell matching and label transfer across sets.

**Supplementary Note 7. Novel visualization options for enhanced biological interpretation of cell heterogeneity**

Cell-ID provides two novel visualization options for the explorative analysis of single-cell RNA-seq data. First, the dimensionality reduction performed through Multiple Correspondence Analysis provides a simultaneous representation of cells and genes on the same principal axes. Thus, Cell-ID allows to visualize cells in the MCA principal components (analogous to a PCA representation), but also to map key gene markers together with the cell representation. In such biplots, multiple gene markers can be displayed at once in a way that, the closer a marker is represented to a given cell, the more specific to them it is. This is illustrated for two independent datasets corresponding to human pancreatic cells ^7^ and mouse airway epithelial cells ^10^, where simultaneous projection of prototypical markers allows to rapidly identify the cell identity of the corresponding regions (**Supplementary Figure 14 A-D**).

Second, Cell-ID provides functional enrichment analysis of the gene signatures obtained for each cell in a dataset, using Gene Ontology terms and pathway annotation databases such KEGG, Reactome and WikiPathways. Functional enrichment analysis may help on the functional interpretation of cell heterogeneity and assist on cell type identification, as illustrated for Schwann cells (**Supplementary Note 3**) and Solitary Chemosensory Cells (SCCs, **Supplementary Note 4**) in the exploratory analysis of pancreas and airway epithelium single-cell RNA-seq datasets, respectively. Thus, functional enrichments can be visualized in a low-dimensionality representation of cells by coloring each cell with an intensity proportional to its −log_10_ p-value for a query functional term or biological pathway (**Supplementary Figure 14 E-F**). Cell-ID R package provides per-cell functional enrichment analysis as a built-in function, which stores the enrichment outputs as additional cell attributes in standard R single-cell data structures, such as SingleCellExperiment and Seurat objects. Such seamless integration allows other single-cell RNA-seq tools to use enrichment scores in alternative cell visualizations, e.g. UMAP^34^ or cell diffusion maps^35^.

## Supplementary Figures

**Supplementary Figure 1.**
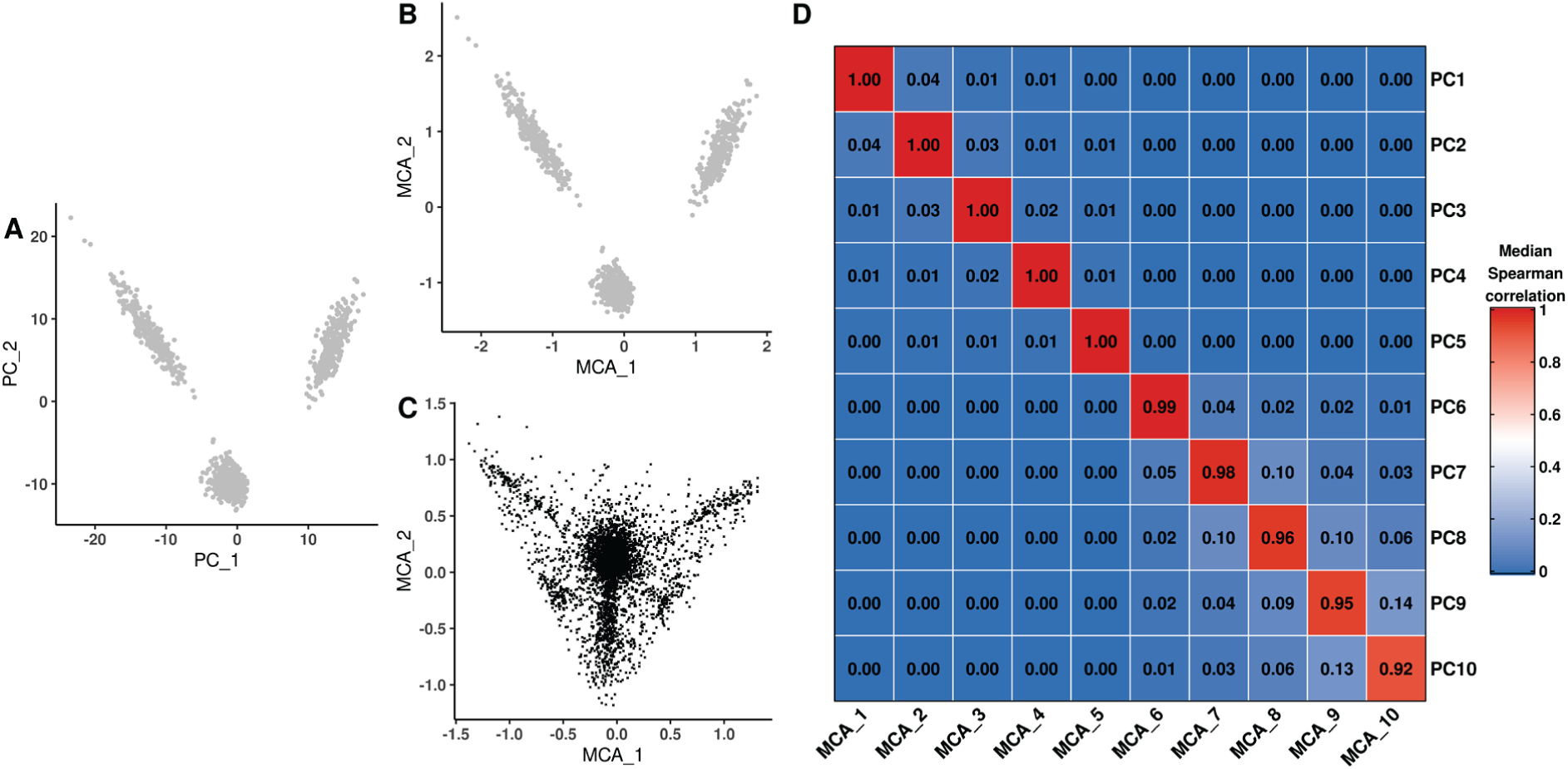
Correspondence between PCA and MCA low-dimensional representations of cells. Low-dimensional representation of a simulated scRNA-seq dataset on the two first principal axes obtained through PCA (**A**, cell projection) or MCA (**B**, cell projection, and **C**, gene projection; **Supplementary Note 1**). (**D**) Median values of Spearman’s correlation coefficient for the relationship between cell coordinates on the first 10 principal axes from MCA (*x*-axis) and the first 10 principal axes from PCA (*y*-axis) over 100 simulated scRNA-seq datasets. Median absolute Spearman’s correlation coefficient values are represented on a color scale ranging from 0 (dark blue) to 1 (dark red), and the precise value is indicated inside.

**Supplementary Figure 2.**
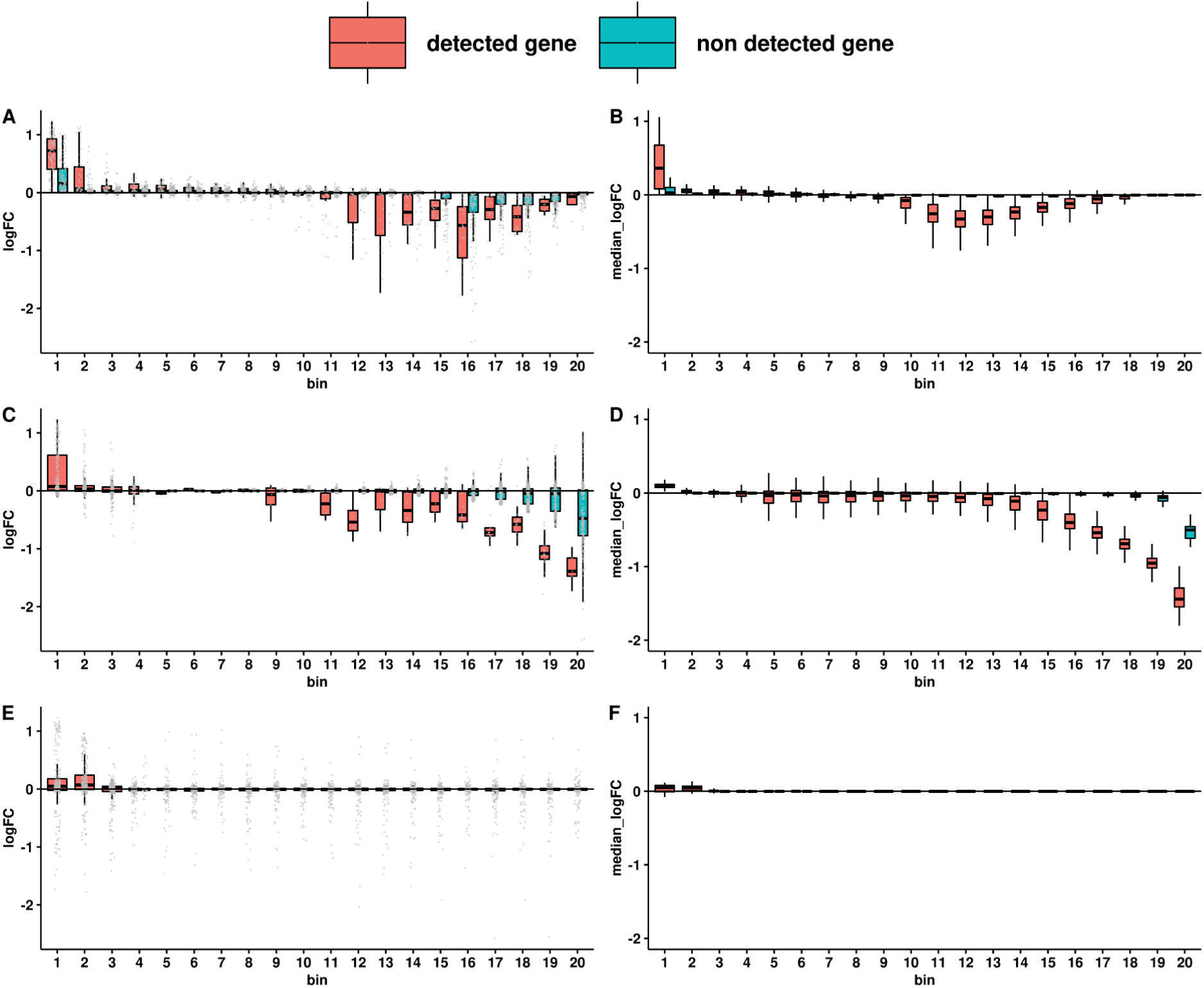
Consistency of the per-cell gene rankings with the gene expression levels of neighboring cells in the MCA space. **(A)** An individual cell was randomly selected from a dataset randomly selected from 100 simulated scRNA-seq sets, each containing 1000 cells and 5000 genes (**Supplementary Note 1**). For each gene (grey dot), this figure shows the log-fold change in expression relative to (i) its mean expression value in the 5% of cells closest to the target cell (*n*=50), and (ii) its mean expression value in the rest of the cells in the dataset (*y*-axis). Genes were grouped into 20 bins of equal size (*x*-axis, on the basis of their ranking relative to the target cell on the MCA space (**Figure 1**). High ranks (on the left side of the *x*-axis) correspond to small gene-to-cell distances, indicating that gene expression is highly specific to the target cell, whereas lower ranks (on the right) reflect a lack of specificity to the target cell. The gene expression values in the target cell were not considered in the assessment of such log-fold changes. For each bin, two boxplots are shown, summarizing (i) the distribution of log-fold changes in expression across genes for which the target cell presented a non-zero count (red); and (ii) the distribution of log-fold changes in expression across genes for which the target cell presented a zero count (blue). **(B)** Analogous figure to **(A)**, showing the generalization of patterns to all individual cells in a randomly selected dataset. Each individual cell was first independently assessed as in (A), and its per-bin median values were extracted to plot a distribution across cells (boxplots in B). Figures analogous to (**A**) and (**B**) are shown when the per-cell gene rankings displayed on the *x*-axis through bins of equal size were obtained from (i) a naïve approach based on the log-fold changes in gene expression observed in a cell relative to all the other cells in the dataset (**C, D**) or (ii) from highest-to-lowest expression values within a cell, with random ranks for ties, as previously described (AUCell, **E, F**).

**Supplementary Figure 3.**
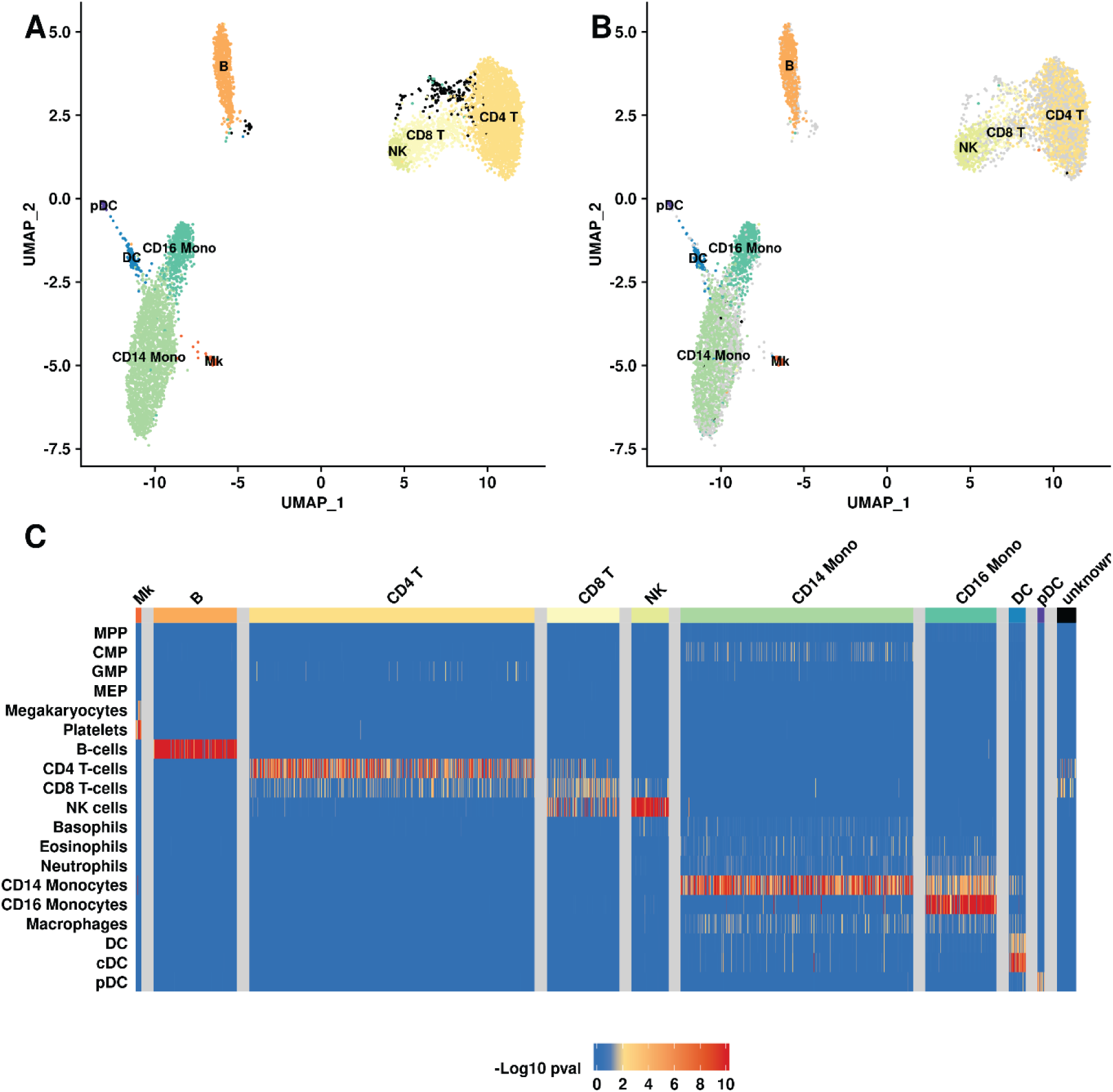
Automatic Cell-ID cell-type identification of human peripheral blood mononuclear cells with pre-established signatures. UMAP representation of 7488 peripheral blood mononuclear cells (PBMCs) profiled with a REAP-seq protocol, with color-coding of the cells according to blood cell type classification based on **(A)** single-cell protein marker levels as provided, and **(B)** Cell-ID predictions based on a reference collection of well-established blood cell signatures. **(C)** Heatmap representing, for each individual cell (displayed in columns), the −log10 transformed *p* value obtained by Cell-ID testing of the association of the gene signature with each of the evaluated pre-established signatures (displayed in rows). The heatmap color scale extends from dark blue (*p* value=1) to yellow (*p* value = 10^−2^) to dark red (*p* value = 10^−10^), with *p* values<10^−10^ fixed at this value). Non-significant associations (*p* value>10^−2^ after Benjamini Hochberg correction for the number of gene signatures tested) are shown in blue. The columns in the heatmaps were grouped by the reference cell-type label, as indicated by the colored bands at the top and in the associated legend.

**Supplementary Figure 4.**
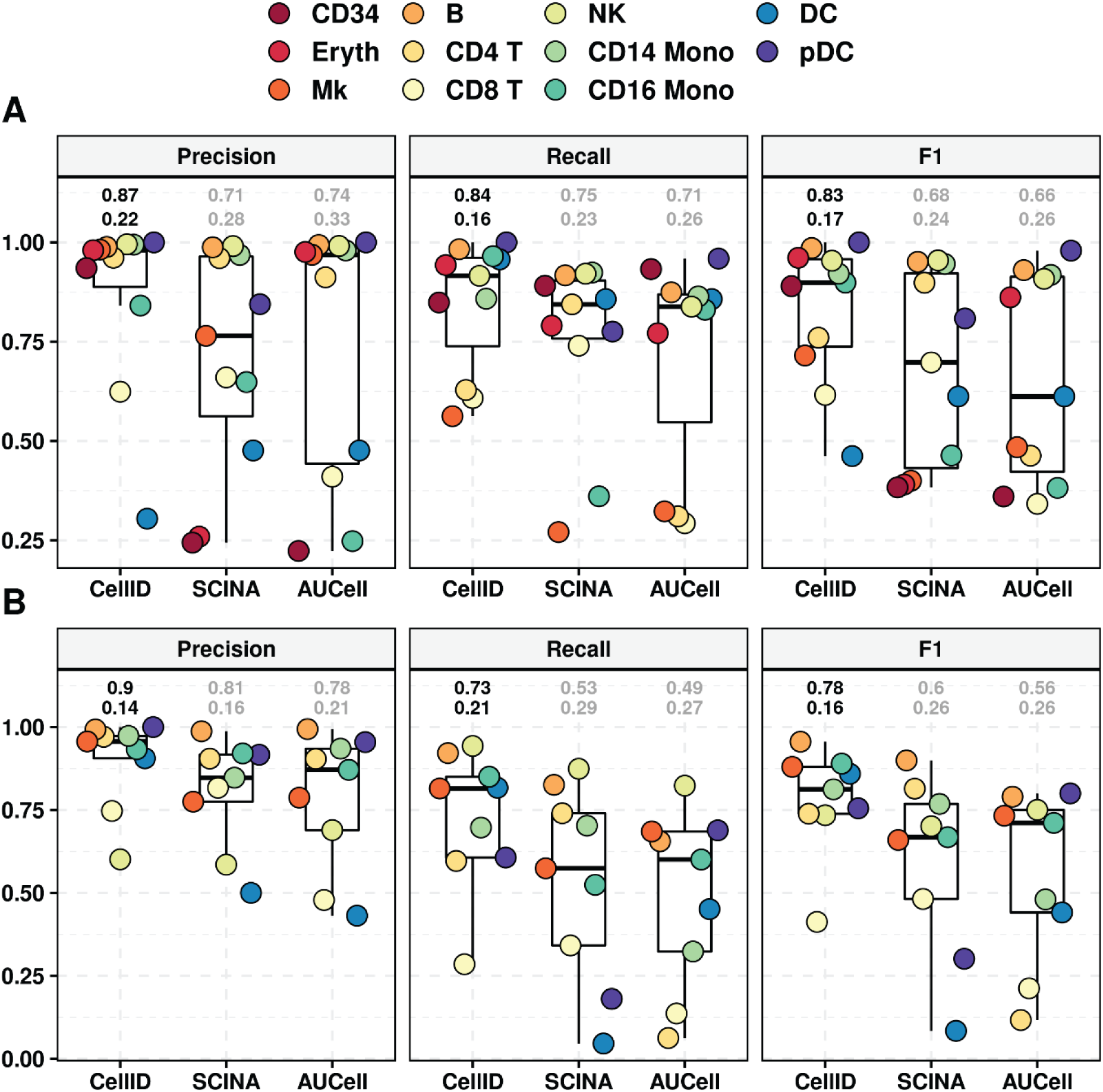
Performance measured through Precision, recall, and F1 scores achieved by Cell-ID, AUCell and SCINA cell type predictions on (**A**) CBMCs Cite-Seq data, and (B) PBMCs Reap-Seq data, for each of the blood cell types reported in the original publications. Boxplots summarize the corresponding scores for each method. The numbers above boxplots denote the global performance (macro F1 score, upper digits) and its standard deviation (lower digits), where the maximum and minimum values across methods are colored in black and grey, respectively.

**Supplementary Figure 5.**
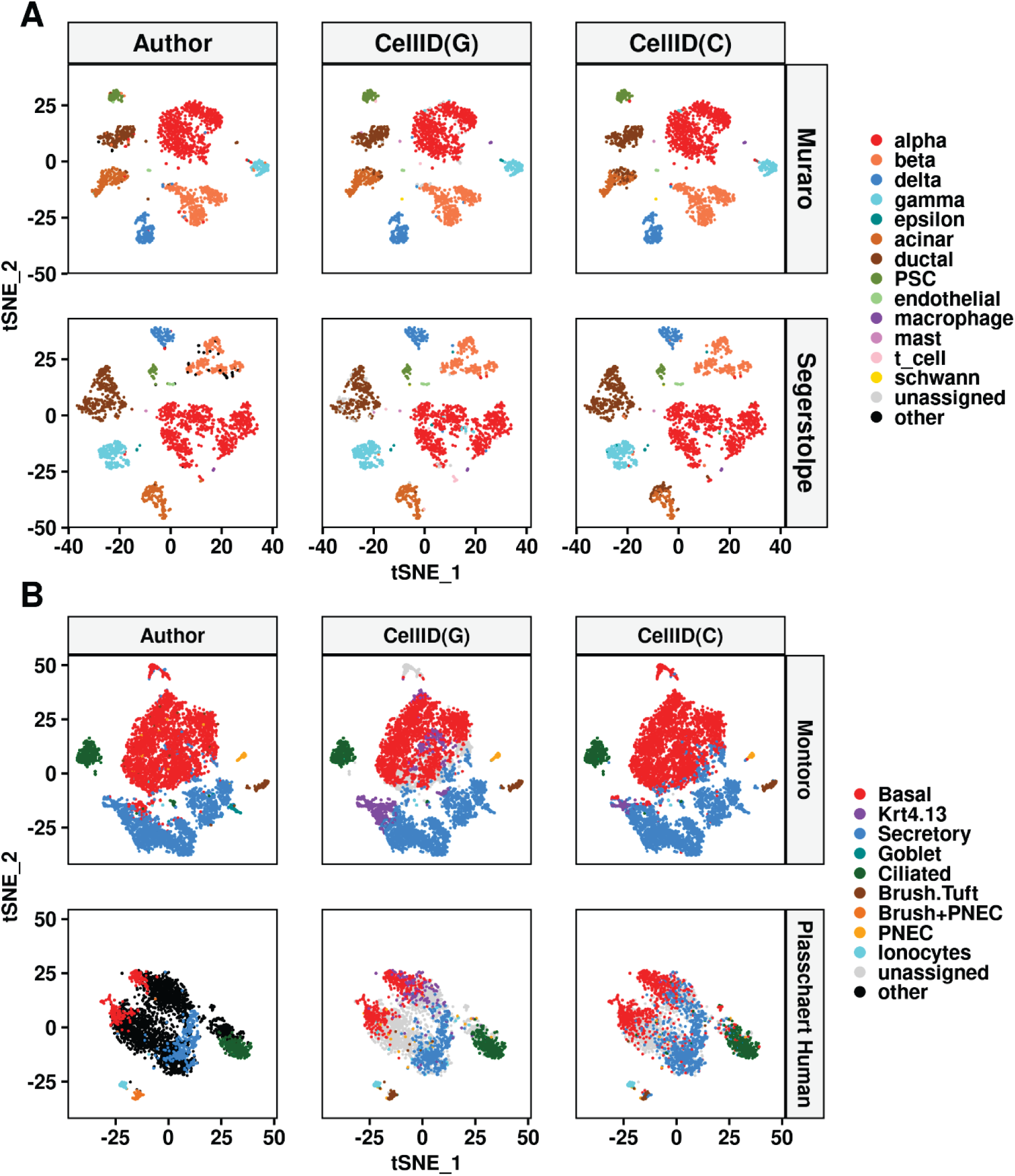
tSNE representation of (**A)** human pancreatic islet cells from the Muraro (top panels) and Segerstolpe (bottom panels) datasets, and (**B)** airway epithelium cells from the Montoro’s mouse dataset (top panels) and Plasschaert’s human dataset (bottom panels). Cells are color-coded according to the cell type labels annotated in the original publications (left panels), as well as by the cell type predictions from Cell-ID(g) (middle panels), and CellID(c) (right panels).

**Supplementary Figure 6.**
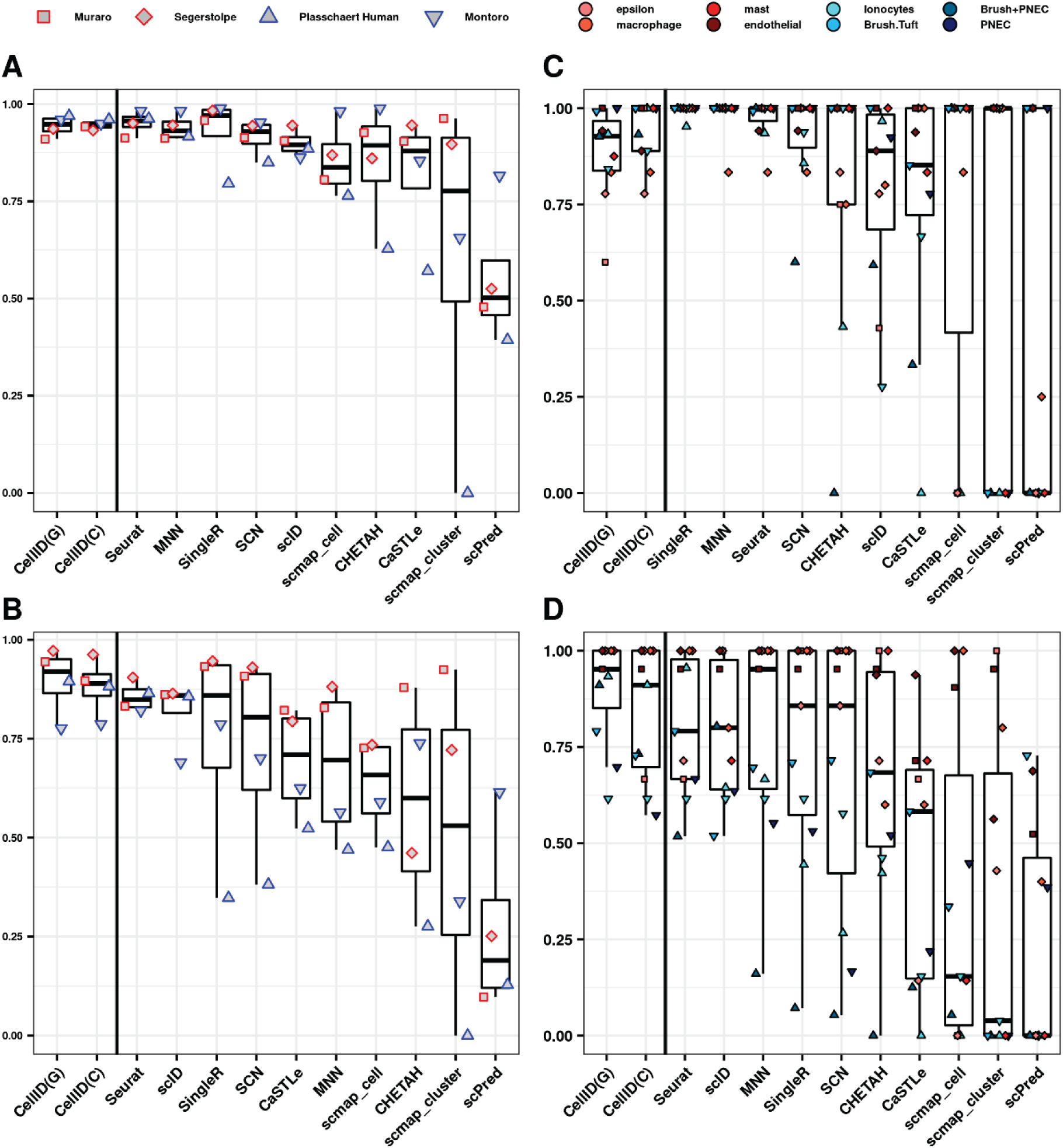
Precision and recall of Cell-ID cell-to-cell matching across independent scRNA-seq datasets from the same or different tissue of origin, within and across species. Performance achieved by Cell-ID(g), Cell-ID(c) and 10 alternative state-of-the-art methods (x-axis), measured through Precision **(A and C)**, and Recall **(B and D)**. Panels are analogous to the ones described in **Figure 3 (A-B)** legend.

**Supplementary Figure 7.**
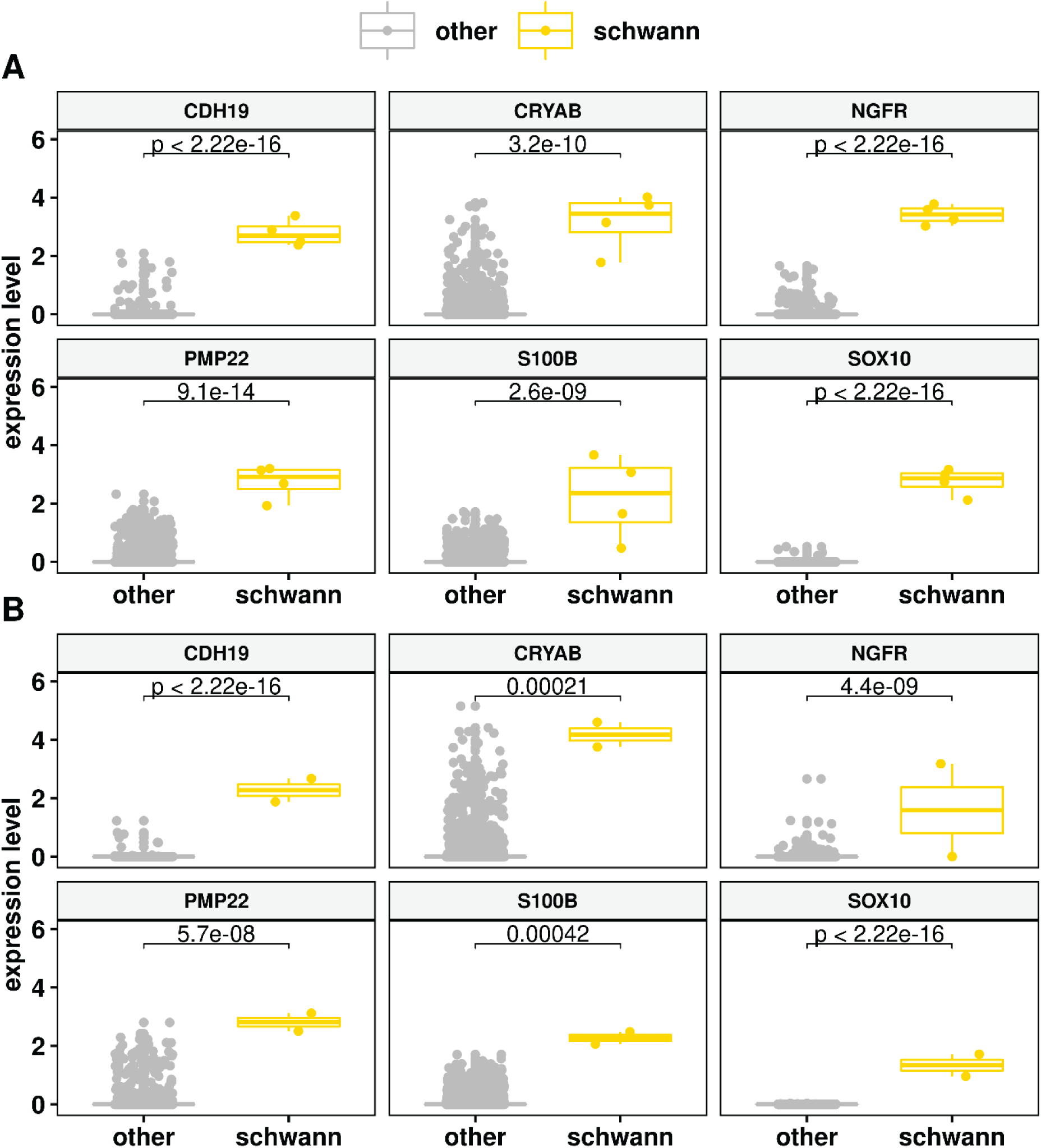
Distribution of log-transformed normalized gene expression levels (y-axis) of Schwann cell markers (SOX10, S100B, CRYAB, NGFR, CDH19, and PMP22) across pancreatic islet cells from (**A)** Segerstolpe and (**B**) Muraro datasets. Values their associated boxplots are represented in yellow for the *n*=4 and n=2 cells in the Muraro and Segerstolpe datasets respectively that where identified by both Cell-ID(c) and Cell-ID(g) as putative Schwann cells, and in grey for the rest of cells. Brackets and *p* values indicate the results of two-sided Wilcoxon rank sum tests comparing the distribution across putative Schwann cells versus the other cells in the dataset.

**Supplementary Figure 8.**
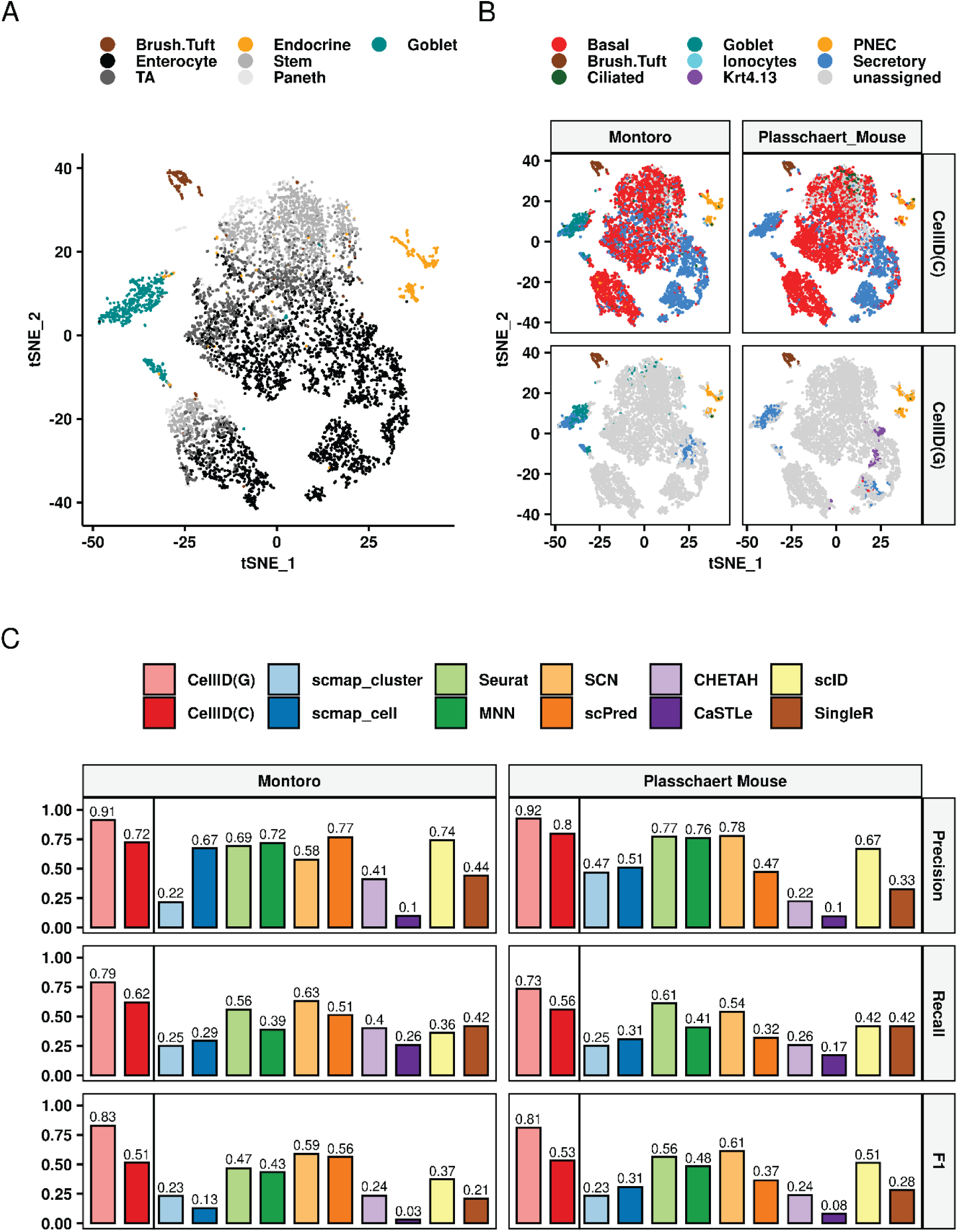
t-SNE representation of 7216 cells from mouse small intestinal epithelium, where dots representing cells are color coded according to (**A**) manual cell type annotations provided in Haber et al. and (**B**) Cell-ID(c) and Cell-ID(g) cell type predictions (top and bottom panels, respectively), using as a reference the mouse airway epithelial gene signatures extracted from Montoro and Plasschaert datasets (left and right panels, respectively). Cells with no significant enrichments were left unassigned and are displayed in grey. (**C**) Precision, Recall and F1 score (y-axis), achieved by Cell-ID(g), Cell-ID(c) and 10 alternative state-of-the-art methods (x-axis), for the label transferring results depicted in **(B)**.

**Supplementary Figure 9.**
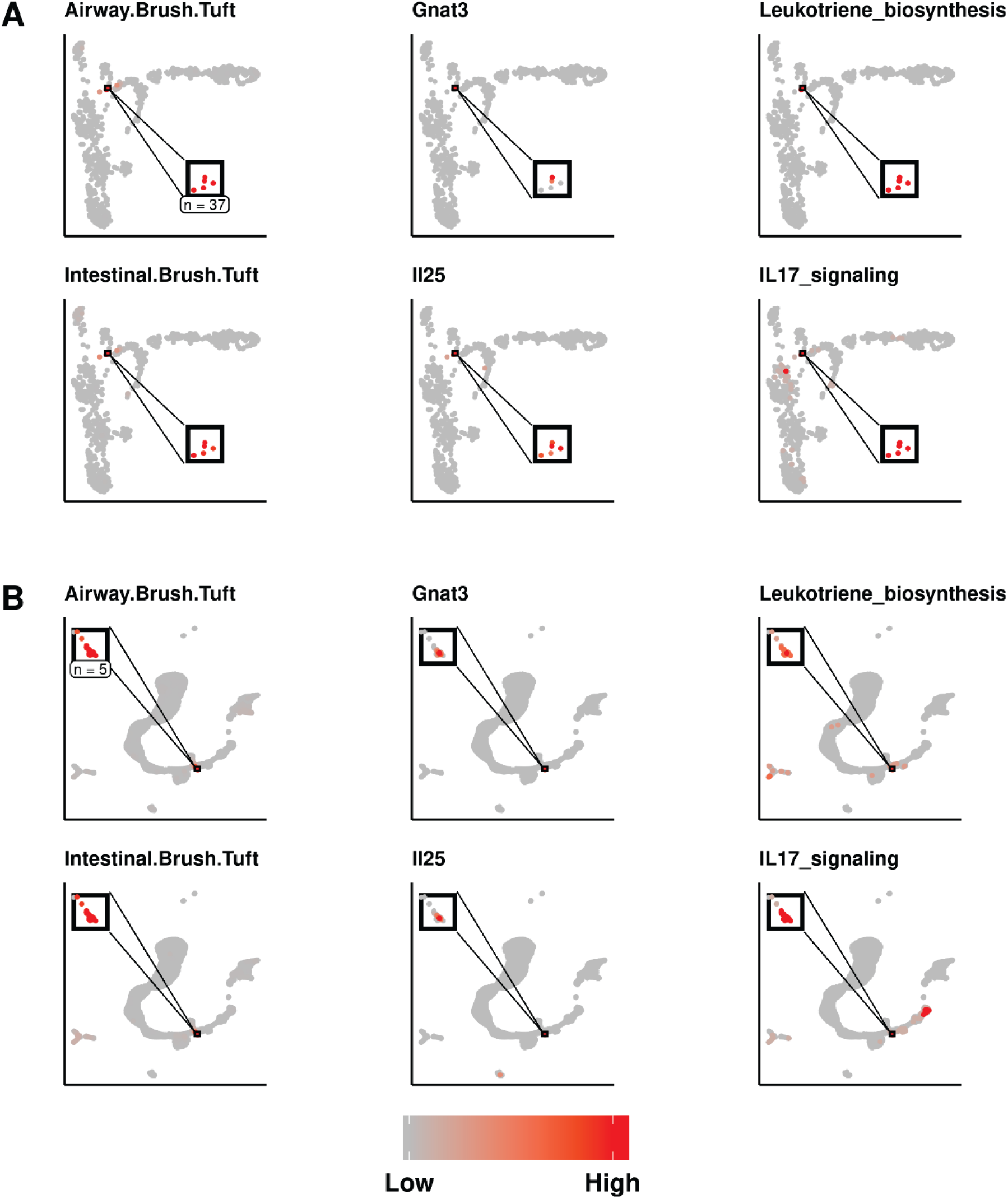
UMAP representation of olfactory epithelium cells from (**A**) Fletcher et al. and (**B**) Wu et al. datasets. The inset panels focus on cells identified by Cell-ID as putative SCCs. Cells represented by dots are color-coded according to (i) their Cell-ID(g) −log_10_ enrichment p-value using as a reference the mouse brush/tuft gene signatures extracted from mouse airway epithelium, and from small intestinal epithelium, as indicated in the panel titles; (ii) the log-normalized expression of two solitary chemosensory cells markers, i.e. interleukin 25 (IL25) and G protein subunit alpha Transducin 3 (GNAT3); and (iii) The −log10 enrichment p-values for 2 functional terms, i.e.: “leukotriene biosynthetic process” (Gene Ontology term GO:0019370) and “interleukin-17-mediated signalling pathway” (GO:0097400). The color scale throughout panels extends from gray (indicating a non-significant *p* value or a lack of detection of gene expression) to red, corresponding to a significant *p* value or high level of gene expression.

**Supplementary Figure 10.**
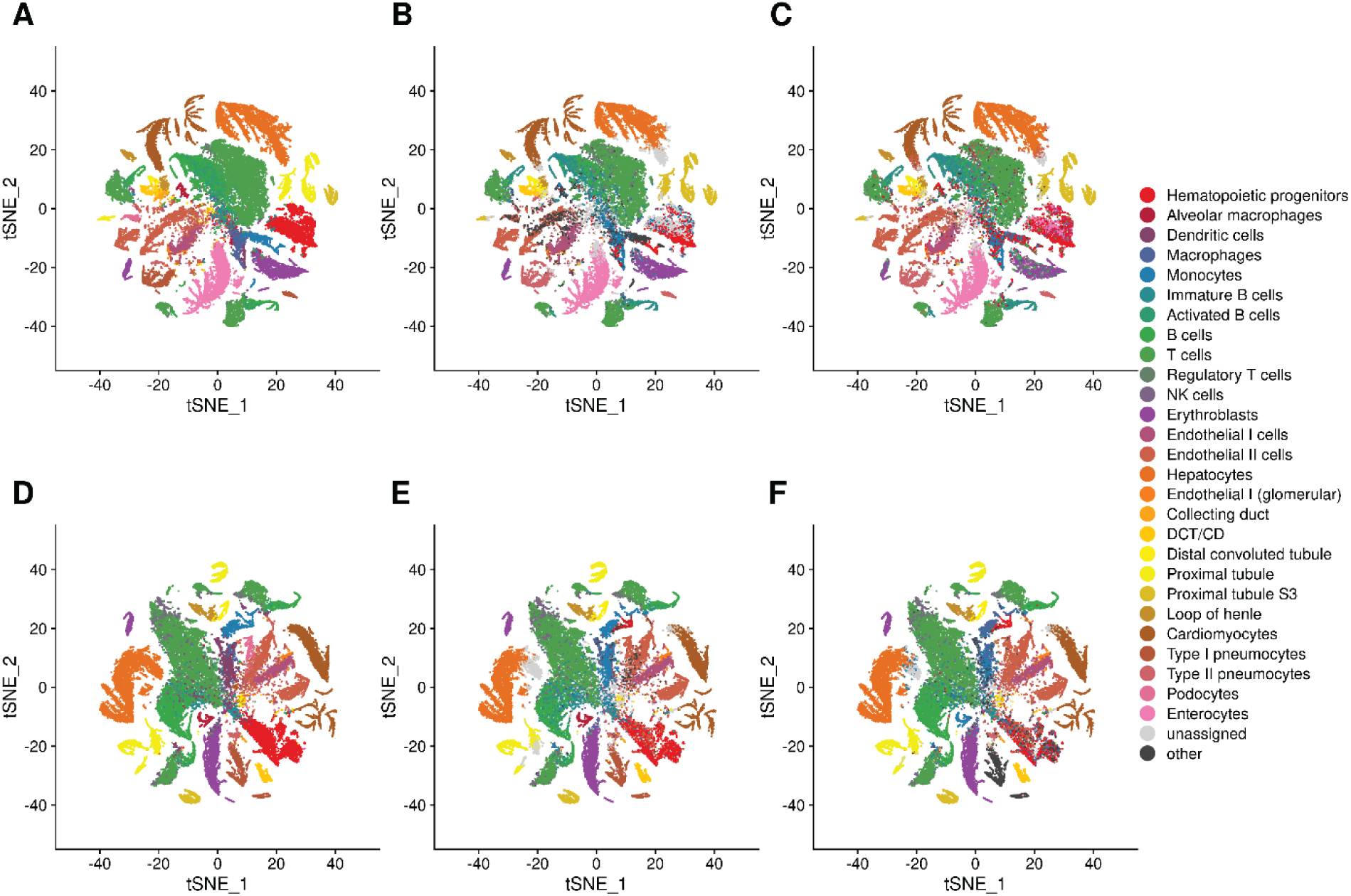
**(A-C)** t-SNE representation of 57370 cells profiled with scATAC-seq from the Mouse ATAC atlas corresponding to the eight tissues for which Tabula Muris SmartSeq scRNA-seq datasets are available, i.e.: heart, kidney, liver, lung, bonne marrow, spleen, thymus and large intestine. **(D-E)** t-SNE representation of 50284 cells profiled with scATAC-seq from the Mouse ATAC atlas corresponding to the seven tissues for which Tabula Muris 10X Genomics scRNA-seq datasets are available, i.e.: heart, kidney, liver, lung, bonne marrow, spleen, thymus. Cells in are colored according to the manually annotated cell types provided by the atlas, regrouped by the categories represented in the legend and described in **Supplementary Table 5**, (**A** and **D**), as well as by the cell type predictions from Cell-ID(g) **(B** and **E)**, and CellID(c) **(C** and **F)** using the group gene signatures extracted from (**A** and **D**, respectively).

**Supplementary Figure 11.**
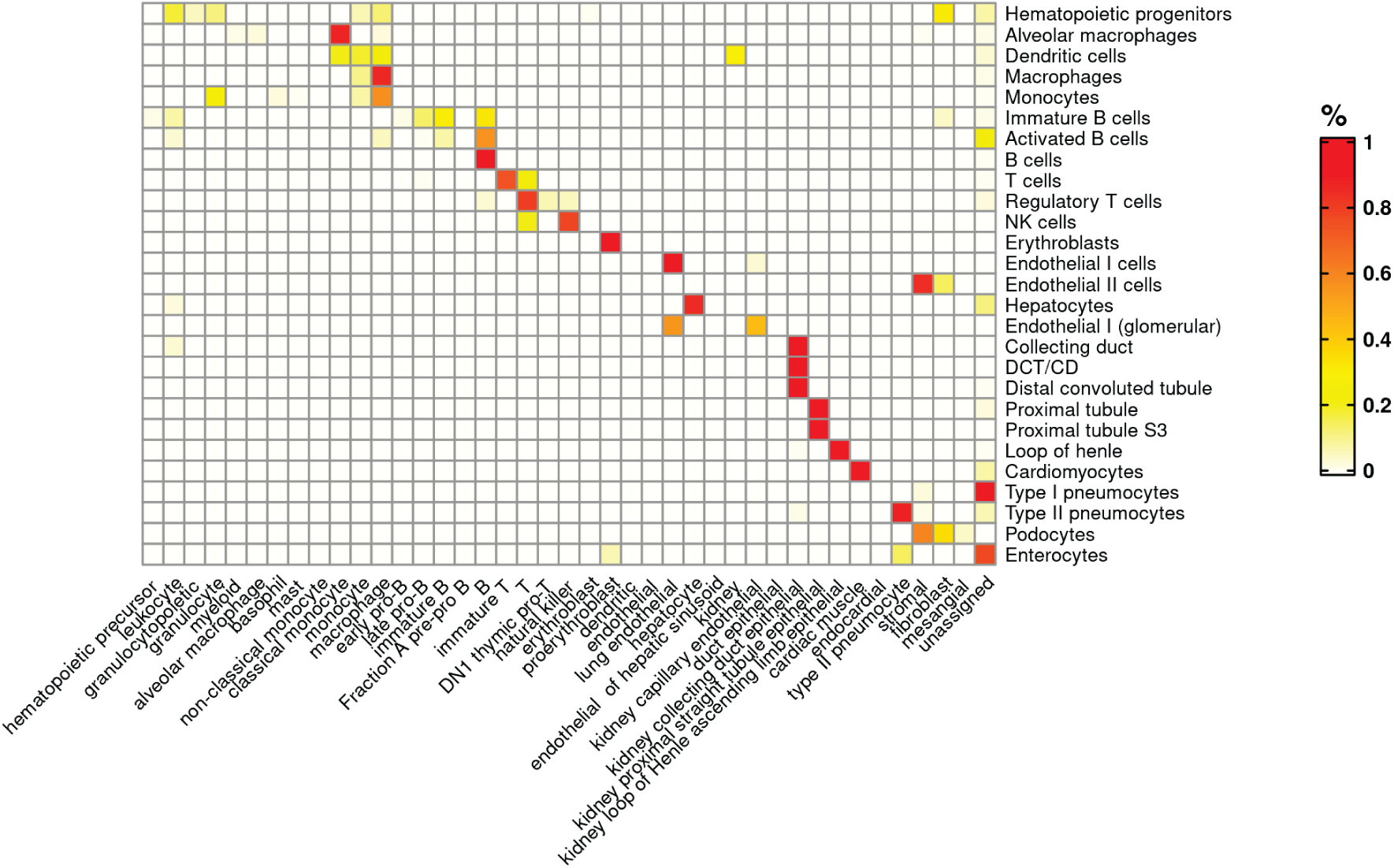
Heatmap representing the confusion matrix between the manually curated cell type annotations from the Mouse ATAC atlas (displayed per rows) and the Cell-ID(c) cell type predictions (displayed per columns) using the gene-signatures extracted from the manually annotated cell types provided by the Tabula Muris mouse cell atlas. The color code in the heatmap represents the ratio *r* of the cell types displayed per rows that are allocated in the cell types represented per columns, ranging from white (*r* = 0) to red (*r* =1).

**Supplementary Figure 12.**
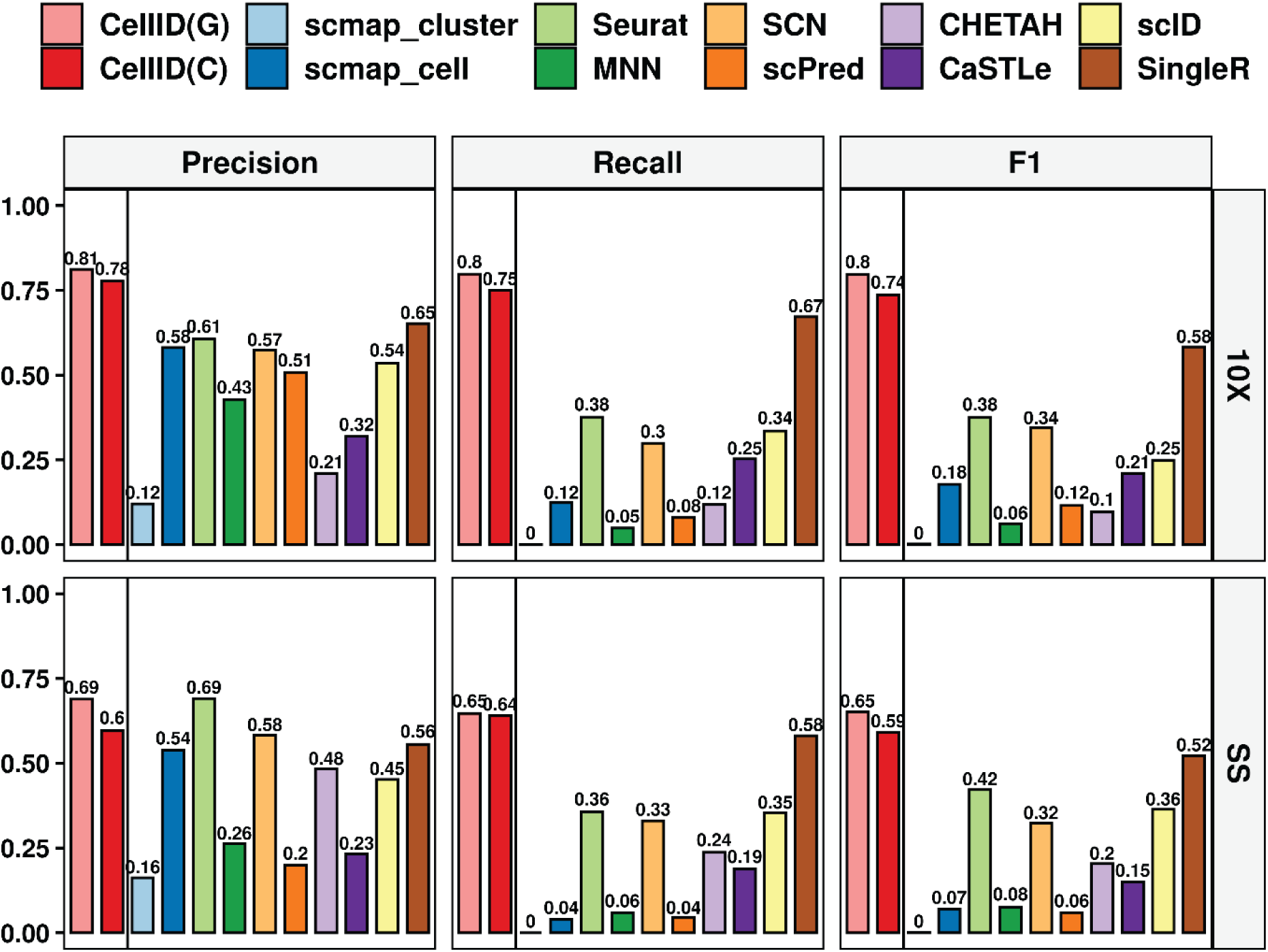
Performance measured as F1 score (y-axis) achieved by Cell-ID(g), Cell-ID(c) and 10 alternative state-of-the-art methods (x-axis), in the cell type label transferring from Tabula Muris scRNAseq to scATAC Atlas for 10X (top panel) and smarts-seq (bottom panel), as depicted in Supplementary Figure 10.

**Supplementary Figure 13.**
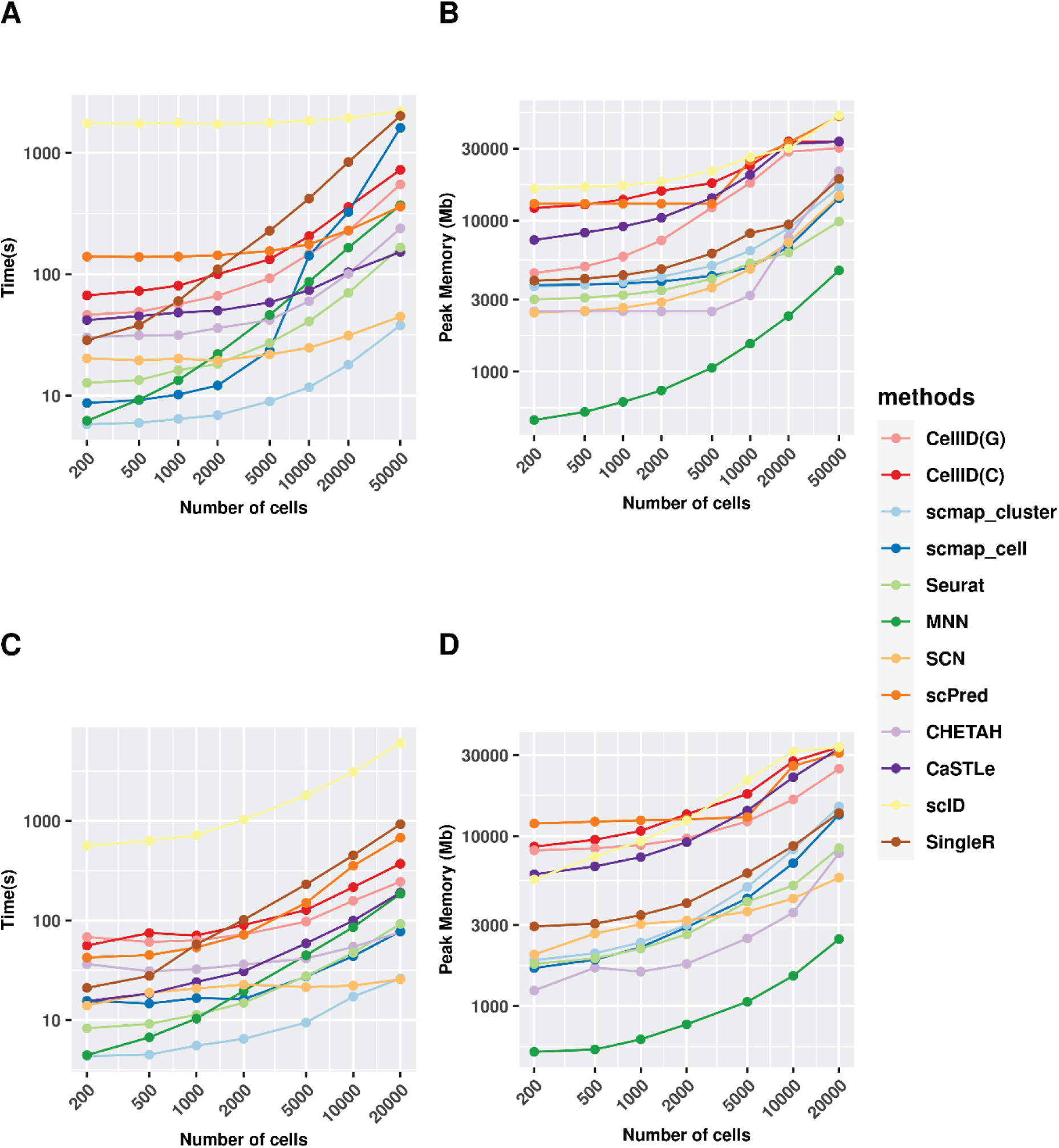
Computational time and memory consumption of label transfer methods on simulated datasets. **(A** and **B)** Line plots showing on the y axis (**A)** the computation time and (**B)** the total memory allocation as a function of the number of cells (*x* axis) randomly sampled from the Tabula Muris 10X data that were used as *query* dataset for cell type label transferring from a *reference* dataset of 5000 randomly sampled cells from the Tabula Muris SmartSeq dataset. Results are plotted for Cell-ID(g), Cell-ID(c) and the 10 state of the art methods evaluated. (**C** and **D**) Line plots showing on the y axis (**C)** the computation time and (**D)** the total memory allocation as a function of the number of cells (*x* axis) randomly sampled from the Tabula Muris SmartSeq data used as *reference* dataset for cell type label transferring on a *query* dataset of 5000 randomly sampled cells from the Tabula Muris 10X data. Results are plotted for Cell-ID(g), Cell-ID(c) and the 10 state of the art methods evaluated.

**Supplementary Figure 14.**
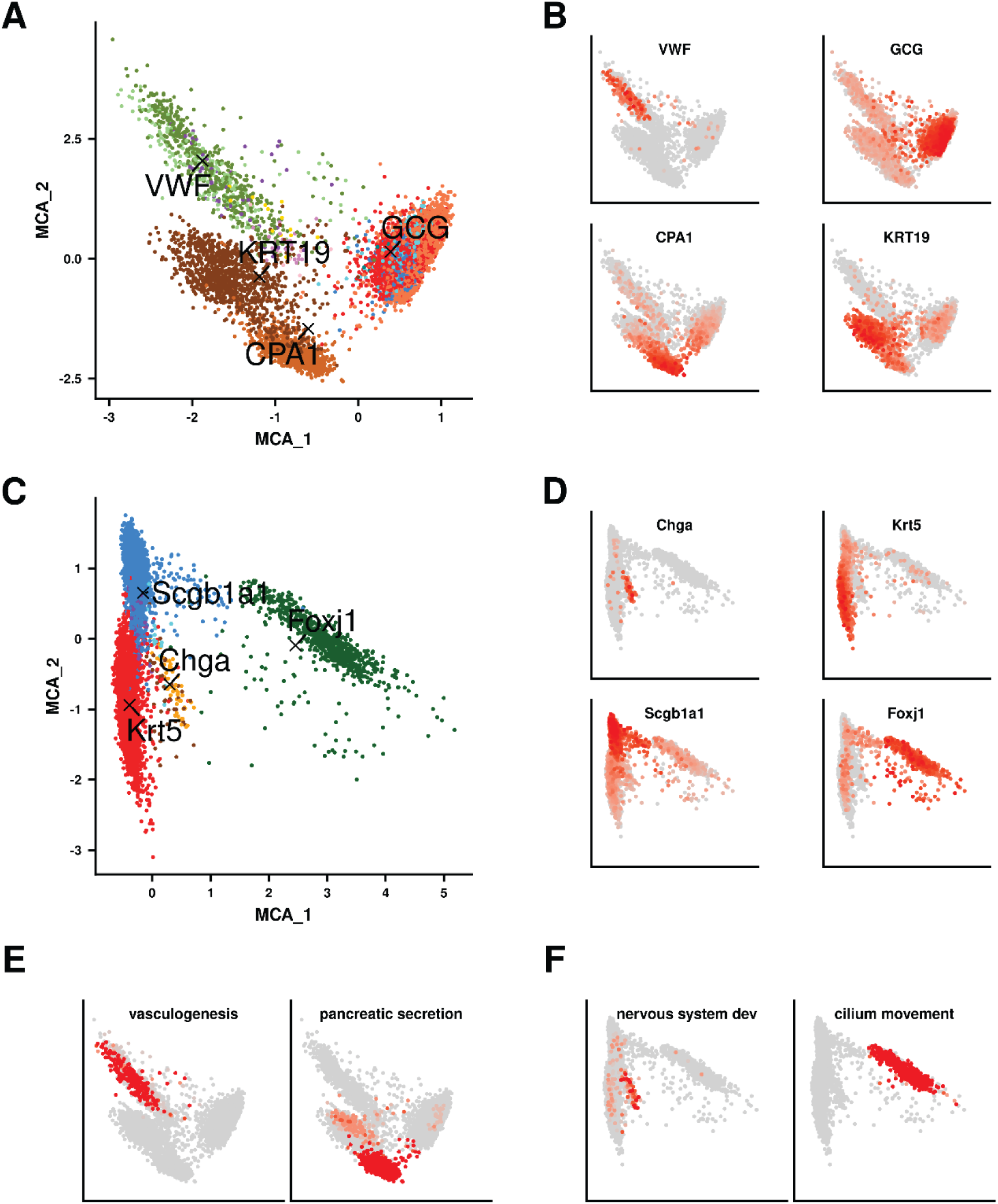
Novel visualization options provided by Cell-ID for the explorative analysis of single-cell RNA-seq data. Simultaneous MCA representation of prototypical marker genes (indicated by back cross symbols and gene name labels) on (**A)** Baron human pancreatic cells, and (**C)** Plasschaert mouse airway cells, color-coded according to the cell type labels annotated in the original publication. **(B)** and **(D)** Equivalent representation of cells in MCA space as depicted in **(A)** and **(C)** where cells are color coded according to their levels of expression for the corresponding selected markers. **(E)** and **(F)** Equivalent representation of cells in MCA space as depicted in **(A)** and **(C)** where cells are color coded according to their functional enrichment −log_10_ p-value for a given functional term or biological pathway, as indicated in the figure titles.

## Supplementary Tables Legends

**Supplementary Table 1**. Functional enrichment analysis results for putative Schwann cells identified in pancreas datasets, and for putative solitary chemosensory cells identified in olfactory epithelium datasets.

**Supplementary Table 2**. Summary of the datasets used in this article.

**Supplementary Table 3**. Summary of external tools and packages used in this article.

**Supplementary Table 4**. Blood cell-type gene markers from the XCell repository used in this study.

**Supplementary Table 5**. Mapping scheme for the original cell population labels from query data to cell population labels in the reference data.

**Supplementary Table 6** Summary of the functional pathways used in this article.

**Supplementary Table 7**. Performance metrics for cell type prediction using pre-established marker lists on blood cells from CITE-seq and REAP-seq datasets. The table reports precision, recall and F1 score for CellID, AUCell and SCINA for each of the evaluated cell types as well as for the overall assessment.

**Supplementary Table 8**. Performance metrics for the prediction of rare cell types from pre-established immune signatures on 100 simulated subsets of CITEseq.

**Supplementary Table 9**. Performance metrics for cell type predictions represented in Figure 3A and Supplementary Figure 6 A-B.

**Supplementary Table 10**. Performance metrics for cell type predictions represented in Figure 3D and Supplementary Figure 8C.

**Supplementary Table 11**. Performance metrics for cell type predictions represented in Figure 4D and Supplementary Figure 12.

## References

1. Teichmann, S. et al. The Human Cell Atlas. eLife 6, (2017).

2. The Human BioMolecular Atlas Program - HuBMAP | NIH Common Fund. https://commonfund.nih.gov/HuBMAP.

3. The LifeTime Initiative. LifeTime FET Flagship https://lifetime-fetflagship.eu/.

4. Lähnemann, D. et al. Eleven grand challenges in single-cell data science. Genome Biol. 21, 31 (2020).

5. Sun, S., Zhu, J., Ma, Y. & Zhou, X. Accuracy, robustness and scalability of dimensionality reduction methods for single-cell RNA-seq analysis. Genome Biol. 20, 269 (2019).

6. Becht, E. et al. Dimensionality reduction for visualizing single-cell data using UMAP. Nat. Biotechnol. 37, 38–44 (2019).

7. Kiselev, V. Y., Andrews, T. S. & Hemberg, M. Challenges in unsupervised clustering of single-cell RNA-seq data. Nat. Rev. Genet. 20, 273–282 (2019).

8. Greenacre, M. J. Theory and applications of correspondence analysis. (1984).

9. Multiple correspondence analysis and related methods. Selected papers based on the presentations at the international conference (CARME 2003), Barcelona, Spain, 29 June to 2 July 2003. Chapman & Hall/CRC Statistics in the Social and Behavioral Sciences Series (Chapman & Hall/CRC, 2006).

10. Aşan, Z. & Greenacre, M. Biplots of fuzzy coded data. Fuzzy Sets Syst. 183, 57–71 (2011).

11. Aibar, S. et al. SCENIC: Single-cell regulatory network inference and clustering. Nat. Methods 14, 1083–1086 (2017).

12. Zhang et al. SCINA: Semi-Supervised Analysis of Single Cells in Silico. Genes 10, 531–531 (2019).

13. Aran, D., Hu, Z. & Butte, A. J. xCell: digitally portraying the tissue cellular heterogeneity landscape. Genome Biol. 18, 220 (2017).

14. Stoeckius, M. et al. Simultaneous epitope and transcriptome measurement in single cells. Nat. Methods 14, 865–868 (2017).

15. Peterson, V. M. et al. Multiplexed quantification of proteins and transcripts in single cells. Nat. Biotechnol. 35, 936–939 (2017).

16. Baron, M. et al. A Single-Cell Transcriptomic Map of the Human and Mouse Pancreas Reveals Inter- and Intra-cell Population Structure. Cell Syst. 3, 346–360 (2016).

17. Segerstolpe, Å. et al. Single-Cell Transcriptome Profiling of Human Pancreatic Islets in Health and Type 2 Diabetes. Cell Metab. 24, 593–607 (2016).

18. Muraro, M. J. et al. A Single-Cell Transcriptome Atlas of the Human Pancreas. Cell Syst. 3, 385–394.e3 (2016).

19. Plasschaert, L. W. et al. A single-cell atlas of the airway epithelium reveals the CFTR-rich pulmonary ionocyte. Nature 560, 377–381 (2018).

20. Montoro, D. T. et al. A revised airway epithelial hierarchy includes CFTR-expressing ionocytes. Orit Rozenblatt-Rosen 4, 19–19 (2018).

21. Kiselev, V. Y., Yiu, A. & Hemberg, M. scmap: projection of single-cell rna-seq data across data sets. 15, 359–359 (2018).

22. Haghverdi, L., Lun, A. T. L., Morgan, M. D. & Marioni, J. C. Batch effects in single-cell RNA-sequencing data are corrected by matching mutual nearest neighbors. Nat. Biotechnol. 36, 421–427 (2018).

23. De Kanter, J. K., Lijnzaad, P., Candelli, T., Margaritis, T. & Holstege, F. C. P. CHETAH: a selective, hierarchical cell type identification method for single-cell RNA sequencing. Nucleic Acids Res. 47, (2019).

24. Lieberman, Y., Rokach, L. & Shay, T. CaSTLe – Classification of single cells by transfer learning: Harnessing the power of publicly available single cell RNA sequencing experiments to annotate new experiments. PLOS ONE 13, e0205499–e0205499 (2018).

25. Boufea, K., Seth, S. & Batada, N. N. scID Uses Discriminant Analysis to Identify Transcriptionally Equivalent Cell Types across Single-Cell RNA-Seq Data with Batch Effect. iScience 23, 100914 (2020).

26. Tan, Y. & Cahan, P. SingleCellNet: A Computational Tool to Classify Single Cell RNA-Seq Data Across Platforms and Across Species. Cell Syst. 9, 207–213.e2 (2019).

27. Alquicira-Hernandez, J., Sathe, A., Ji, H. P., Nguyen, Q. & Powell, J. E. ScPred: Accurate supervised method for cell-type classification from single-cell RNA-seq data. Genome Biol. 20, 264–264 (2019).

28. Aran, D. et al. Reference-based analysis of lung single-cell sequencing reveals a transitional profibrotic macrophage. Nat. Immunol. 20,.

29. Stuart, T. et al. Comprehensive Integration of Single-Cell Data. Cell 177, 1888–1902.e21 (2019).

30. Haber, A. L. et al. A single-cell survey of the small intestinal epithelium. (2017) doi: 10.1038/nature24489.

31. Wu, Y. et al. A Population of Navigator Neurons Is Essential for Olfactory Map Formation during the Critical Period Article A Population of Navigator Neurons Is Essential for Olfactory Map Formation during the Critical Period. Neuron 100, 1066–1082.e6 (2018).

32. Fletcher, R. B. et al. Deconstructing Olfactory Stem Cell Trajectories at Single-Cell Resolution. Cell Stem Cell 20, 817–830.e8 (2017).

33. Ualiyeva, S. et al. Airway brush cells generate cysteinyl leukotrienes through the ATP sensor P2Y2. Sci. Immunol. 5, eaax7224–eaax7224 (2020).

34. Bankova, L. G. et al. The cysteinyl leukotriene 3 receptor regulates expansion of IL-25–producing airway brush cells leading to type 2 inflammation. Sci. Immunol. 3, (2018).

35. Schaum, N. et al. Single-cell transcriptomics of 20 mouse organs creates a Tabula Muris the tabula Muris consortium*. (2018) doi: 10.1038/s41586-018-0590-4.

36. Cusanovich, D. A. et al. A Single-Cell Atlas of In Vivo Mammalian Chromatin Accessibility. Cell 174, 1309–1324.e18 (2018).

37. Franzén, O., Gan, L.-M. & Björkegren, J. L. M. PanglaoDB: a web server for exploration of mouse and human single-cell RNA sequencing data. Database 2019, (2019).

38. Zhang, X. et al. CellMarker: a manually curated resource of cell markers in human and mouse. Nucleic Acids Res. 47, D721–D728 (2019).

39. Liberzon, A. et al. The Molecular Signatures Database Hallmark Gene Set Collection. Cell Syst. 1, 417–425 (2015).

40. The Gene Ontology (GO) database and informatics resource. Nucleic Acids Res. 32, D258–D261 (2004).

41. Jassal, B. et al. The reactome pathway knowledgebase. Nucleic Acids Res. 48, D498–D503 (2020).

42. Kanehisa, M., Sato, Y., Kawashima, M., Furumichi, M. & Tanabe, M. KEGG as a reference resource for gene and protein annotation. Nucleic Acids Res. 44, 457–462 (2015).

43. Slenter, D. N. et al. WikiPathways: a multifaceted pathway database bridging metabolomics to other omics research. Nucleic Acids Res. 46, D661–D667 (2018).

## Methods references

1. Stoeckius, M. et al. Simultaneous epitope and transcriptome measurement in single cells. Nat. Methods 14, 865–868 (2017).

2. Peterson, V. M. et al. Multiplexed quantification of proteins and transcripts in single cells. Nat. Biotechnol. 35, 936–939 (2017).

3. Stuart, T. et al. Comprehensive Integration of Single-Cell Data. Cell 177, 1888–1902.e21 (2019).

4. Baron, M. et al. A Single-Cell Transcriptomic Map of the Human and Mouse Pancreas Reveals Inter- and Intra-cell Population Structure. Cell Syst. 3, 346–360 (2016).

5. Muraro, M. J. et al. A Single-Cell Transcriptome Atlas of the Human Pancreas. Cell Syst. 3, 385–394.e3 (2016).

6. Segerstolpe, Å. et al. Single-Cell Transcriptome Profiling of Human Pancreatic Islets in Health and Type 2 Diabetes. Cell Metab. 24, 593–607 (2016).

7. scRNAseq. Bioconductor http://bioconductor.org/packages/scRNAseq/.

8. Plasschaert, L. W. et al. A single-cell atlas of the airway epithelium reveals the CFTR-rich pulmonary ionocyte. Nature 560, 377–381 (2018).

9. Montoro, D. T. et al. A revised airway epithelial hierarchy includes CFTR-expressing ionocytes. Orit Rozenblatt-Rosen 4, 19–19 (2018).

10. Haber, A. L. et al. A single-cell survey of the small intestinal epithelium. (2017) doi: 10.1038/nature24489.

11. Fletcher, R. B. et al. Deconstructing Olfactory Stem Cell Trajectories at Single-Cell Resolution. Cell Stem Cell 20, 817–830.e8 (2017).

12. Wu, Y. et al. A Population of Navigator Neurons Is Essential for Olfactory Map Formation during the Critical Period Article A Population of Navigator Neurons Is Essential for Olfactory Map Formation during the Critical Period. Neuron 100, 1066–1082.e6 (2018).

13. Schaum, N. et al. Single-cell transcriptomics of 20 mouse organs creates a Tabula Muris the tabula Muris consortium*. (2018) doi: 10.1038/s41586-018-0590-4.

14. Cusanovich, D. A. et al. A Single-Cell Atlas of In Vivo Mammalian Chromatin Accessibility. Cell 174, 1309–1324.e18 (2018).

15. Zerbino, D. R. et al. Ensembl 2018. Nucleic Acids Res. 46, D754–D761 (2018).

16. Durinck, S., Spellman, P. T., Birney, E. & Huber, W. Mapping Identifiers for the Integration of Genomic Datasets with the R/Bioconductor package biomaRt. Nat. Protoc. 4, 1184–1191 (2009).

17. Multiple correspondence analysis and related methods. Selected papers based on the presentations at the international conference (CARME 2003), Barcelona, Spain, 29 June to 2 July 2003. Chapman & Hall/CRC Statistics in the Social and Behavioral Sciences Series (Chapman & Hall/CRC, 2006).

18. Greenacre, M. J. Theory and applications of correspondence analysis. (1984).

19. Lebart, L., Morineau, A. & Warwick, K. M. Multivariate descriptive statistical analysis. Correspondence analysis and related techniques for large matrices. Transl. from French by Elisabeth Moraillon Berry. With a foreword by Herman P. Friedman. Wiley Series in Probability and Mathematical Statistics (John Wiley & Sons, Hoboken, NJ, 1984).

20. Aşan, Z. & Greenacre, M. Biplots of fuzzy coded data. Fuzzy Sets Syst. 183, 57–71 (2011).

21. Greenacre. Chapter 8. in Biplots in practice 79–88 (2010).

22. Benjamini, Y. & Hochberg, Y. Controlling the False Discovery Rate: A Practical and Powerful Approach to Multiple Testing. J. R. Stat. Soc. Ser. B Methodol. 57, 289–300 (1995).

23. Zappia, L., Phipson, B. & Oshlack, A. Splatter: Simulation of single-cell RNA sequencing data. Genome Biol. 18, 174–174 (2017).

24. Zhang et al. SCINA: Semi-Supervised Analysis of Single Cells in Silico. Genes 10, 531–531 (2019).

25. Aibar, S. et al. SCENIC: Single-cell regulatory network inference and clustering. Nat. Methods 14, 1083–1086 (2017).

26. Aran, D., Hu, Z. & Butte, A. J. xCell: digitally portraying the tissue cellular heterogeneity landscape. Genome Biol. 18, 220 (2017).

27. Haghverdi, L., Lun, A. T. L., Morgan, M. D. & Marioni, J. C. Batch effects in single-cell RNA-sequencing data are corrected by matching mutual nearest neighbors. Nat. Biotechnol. 36, 421–427 (2018).

28. Tan, Y. & Cahan, P. SingleCellNet: A Computational Tool to Classify Single Cell RNA-Seq Data Across Platforms and Across Species. Cell Syst. 9, 207–213.e2 (2019).

29. De Kanter, J. K., Lijnzaad, P., Candelli, T., Margaritis, T. & Holstege, F. C. P. CHETAH: a selective, hierarchical cell type identification method for single-cell RNA sequencing. Nucleic Acids Res. 47, (2019).

30. Jassal, B. et al. The reactome pathway knowledgebase. Nucleic Acids Res. 48, D498–D503 (2020).

31. Kanehisa, M., Sato, Y., Kawashima, M., Furumichi, M. & Tanabe, M. KEGG as a reference resource for gene and protein annotation. Nucleic Acids Res. 44, 457–462 (2015).

32. Slenter, D. N. et al. WikiPathways: a multifaceted pathway database bridging metabolomics to other omics research. Nucleic Acids Res. 46, D661–D667 (2018).

33. The Gene Ontology (GO) database and informatics resource. Nucleic Acids Res. 32, D258–D261 (2004).

34. Chen, E. Y. et al. Enrichr: interactive and collaborative HTML5 gene list enrichment analysis tool. BMC Bioinformatics 14, 128 (2013).

35. Becht, E. et al. Dimensionality reduction for visualizing single-cell data using UMAP. Nat. Biotechnol. 37, 38–44 (2019).

36. Kobak, D. & Berens, P. The art of using t-SNE for single-cell transcriptomics. Nat. Commun. 10, 5416 (2019).

## Supplementary references

1. Aibar, S. et al. SCENIC: Single-cell regulatory network inference and clustering. Nat. Methods 14, 1083–1086 (2017).

2. Stoeckius, M. et al. Simultaneous epitope and transcriptome measurement in single cells. Nat. Methods 14, 865–868 (2017).

3. Peterson, V. M. et al. Multiplexed quantification of proteins and transcripts in single cells. Nat. Biotechnol. 35, 936–939 (2017).

4. Aran, D., Hu, Z. & Butte, A. J. xCell: digitally portraying the tissue cellular heterogeneity landscape. Genome Biol. 18, 220 (2017).

5. Zhang et al. SCINA: Semi-Supervised Analysis of Single Cells in Silico. Genes 10, 531–531 (2019).

6. Zhang, A. W. et al. Probabilistic cell-type assignment of single-cell RNA-seq for tumor microenvironment profiling. Nat. Methods 16, 1007–1015 (2019).

7. Baron, M. et al. A Single-Cell Transcriptomic Map of the Human and Mouse Pancreas Reveals Inter- and Intra-cell Population Structure. Cell Syst. 3, 346–360 (2016).

8. Muraro, M. J. et al. A Single-Cell Transcriptome Atlas of the Human Pancreas. Cell Syst. 3, 385–394.e3 (2016).

9. Segerstolpe, Å. et al. Single-Cell Transcriptome Profiling of Human Pancreatic Islets in Health and Type 2 Diabetes. Cell Metab. 24, 593–607 (2016).

10. Plasschaert, L. W. et al. A single-cell atlas of the airway epithelium reveals the CFTR-rich pulmonary ionocyte. Nature 560, 377–381 (2018).

11. Montoro, D. T. et al. A revised airway epithelial hierarchy includes CFTR-expressing ionocytes. Orit Rozenblatt-Rosen 4, 19–19 (2018).

12. Stuart, T. et al. Comprehensive Integration of Single-Cell Data. Cell 177, 1888–1902.e21 (2019).

13. Haghverdi, L., Lun, A. T. L., Morgan, M. D. & Marioni, J. C. Batch effects in single-cell RNA-sequencing data are corrected by matching mutual nearest neighbors. Nat. Biotechnol. 36, 421–427 (2018).

14. Kiselev, V. Y., Yiu, A. & Hemberg, M. scmap: projection of single-cell rna-seq data across data sets. 15, 359–359 (2018).

15. Boufea, K., Seth, S. & Batada, N. N. scID Uses Discriminant Analysis to Identify Transcriptionally Equivalent Cell Types across Single-Cell RNA-Seq Data with Batch Effect. iScience 23, 100914 (2020).

16. Aran, D. et al. Reference-based analysis of lung single-cell sequencing reveals a transitional profibrotic macrophage. Nat. Immunol. 20,.

17. Lieberman, Y., Rokach, L. & Shay, T. CaSTLe – Classification of single cells by transfer learning: Harnessing the power of publicly available single cell RNA sequencing experiments to annotate new experiments. PLOS ONE 13, e0205499–e0205499 (2018).

18. Tan, Y. & Cahan, P. SingleCellNet: A Computational Tool to Classify Single Cell RNA-Seq Data Across Platforms and Across Species. Cell Syst. 9, 207–213.e2 (2019).

19. De Kanter, J. K., Lijnzaad, P., Candelli, T., Margaritis, T. & Holstege, F. C. P. CHETAH: a selective, hierarchical cell type identification method for single-cell RNA sequencing. Nucleic Acids Res. 47, (2019).

20. Bhatheja, K. & Field, J. Schwann cells: Origins and role in axonal maintenance and regeneration. Int. J. Biochem. Cell Biol. 38, 1995–1999 (2006).

21. Chernousov, M. A. et al. Development/Plasticity/Repair Glypican-1 and 4(V) Collagen Are Required for Schwann Cell Myelination. (2006) doi: 10.1523/JNEUROSCI.2544-05.2006.

22. Kanehisa, M., Sato, Y., Kawashima, M., Furumichi, M. & Tanabe, M. KEGG as a reference resource for gene and protein annotation. Nucleic Acids Res. 44, 457–462 (2015).

23. Jassal, B. et al. The reactome pathway knowledgebase. Nucleic Acids Res. 48, D498–D503 (2020).

24. Slenter, D. N. et al. WikiPathways: a multifaceted pathway database bridging metabolomics to other omics research. Nucleic Acids Res. 46, D661–D667 (2018).

25. Bankova, L. G. et al. The cysteinyl leukotriene 3 receptor regulates expansion of IL-25–producing airway brush cells leading to type 2 inflammation. Sci. Immunol. 3, (2018).

26. Ualiyeva, S. et al. Airway brush cells generate cysteinyl leukotrienes through the ATP sensor P2Y2. Sci. Immunol. 5, eaax7224–eaax7224 (2020).

27. Bouchery, T. & Marsland, B. J. Airway brush cells: Not as “tuft” as you might think. Sci. Immunol. 3, (2018).

28. Kohanski, M. A. et al. Solitary chemosensory cells are a primary epithelial source of IL-25 in patients with chronic rhinosinusitis with nasal polyps. J. Allergy Clin. Immunol. 142, 460–469.e7 (2018).

29. Haber, A. L. et al. A single-cell survey of the small intestinal epithelium. (2017) doi: 10.1038/nature24489.

30. Wu, Y. et al. A Population of Navigator Neurons Is Essential for Olfactory Map Formation during the Critical Period Article A Population of Navigator Neurons Is Essential for Olfactory Map Formation during the Critical Period. Neuron 100, 1066–1082.e6 (2018).

31. Fletcher, R. B. et al. Deconstructing Olfactory Stem Cell Trajectories at Single-Cell Resolution. Cell Stem Cell 20, 817–830.e8 (2017).

32. Schaum, N. et al. Single-cell transcriptomics of 20 mouse organs creates a Tabula Muris the tabula Muris consortium*. (2018) doi: 10.1038/s41586-018-0590-4.

33. Cusanovich, D. A. et al. A Single-Cell Atlas of In Vivo Mammalian Chromatin Accessibility. Cell 174, 1309–1324.e18 (2018).

34. Becht, E. et al. Dimensionality reduction for visualizing single-cell data using UMAP. Nat. Biotechnol. 37, 38–44 (2019).

35. Haghverdi, L., Buettner, F. & Theis, F. J. Diffusion maps for high-dimensional single-cell analysis of differentiation data. Bioinformatics 31, 2989–2998 (2015).

